# Pain is so close to pleasure: the same dopamine neurons can mediate approach and avoidance in *Drosophila*

**DOI:** 10.1101/2021.10.04.463010

**Authors:** Daniel Döringer, Christian Rohrsen, Benjamin de Bivort, Wolf Huetteroth, Björn Brembs

## Abstract

Dopaminergic neurons (DANs) have long been implicated in reward, punishment, and value processing across animal phyla, yet whether individual DAN subpopulations convey generalized valence signals or serve more specialized functions remains contested. We addressed this question directly by screening about fifty *Drosophila* DAN subpopulations for their ability to drive consistent approach or avoidance across different operant behavioral paradigms in which flies controlled contingent optogenetic DAN activation through their own behavior. No single DAN subpopulation drove consistent preferences across all paradigms, and DAN activation produced substantially weaker valence effects than activation of sensory neurons used as positive controls. One subpopulation produced a striking time-dependent valence switch: initial aversion gave way to approach over approximately ten minutes of operant engagement. This switch required both dopamine and active behavioral contingency, demonstrating that agency itself is a necessary condition for such a valence switch. Together, our results suggest that individual DANs function as specialized, context-sensitive reporters for specific goals rather than general-purpose value signals, and that their apparent generality under some circumstances may reflect the consequences of non-contingent DAN stimulation rather than a true unified valence code.

## Introduction

*How strange would appear to be this thing that men call pleasure! And how curiously it is related to what is thought to be its opposite, pain! The two will never be found together in a man, and yet if you seek the one and obtain it, you are almost bound always to get the other as well, just as though they were both attached to one and the same head. Wherever the one is found, the other follows up behind. So, in my case, since I had pain in my leg as a result of the fetters, pleasure seems to have come to follow it up*.

> —Plato, Phaedo

How the brain assigns value to sensory experience is a central question in neuroscience. Dopamine neurons (DANs) have long been implicated in this process across animal phyla, and a large vertebrate literature assigns them a central role in reward and punishment. In classical conditioning paradigms, midbrain DANs encode reward prediction errors (Bayer and Glimcher, 2005; Frank, 2025; Montague et al., 1996; Schultz, 2019, 2017, 2016a; Schultz et al., 1993), and optogenetic activation of these neurons can substitute for natural rewards in operant tasks (Kremer et al., 2020). These findings inspired the influential proposal that dopamine implements a “common currency” of value, converting diverse sensory rewards — food, water, sex, money — into a shared neural representation (Berridge and Kringelbach, 2015; Grabenhorst and Rolls, 2011; Kobayashi and Hsu, 2019; Lak et al., 2014; Landreth and Bickle, 2008; Levy and Glimcher, 2012; Matsumoto et al., 2016; Schultz et al., 2015). Within this framework, appetitive outcomes are encoded by one population of DANs, while separate populations encode aversive events (Hu, 2016; Jovanoski and Waddell, 2025; Lammel et al., 2014; Lopez and Lerner, 2025; Pignatelli and Bonci, 2015; Verharen et al., 2020; Warlow et al., 2024; Wenzel et al., 2015).

This concept is now under pressure across animal phyla. Increasing evidence for functional and genetic heterogeneity among dopamine neurons — even within tightly circumscribed anatomical regions — has begun to challenge unified accounts of dopaminergic valuation in vertebrates (e.g., Adam, 2026; Azcorra et al., 2023; Elum et al., 2024; Engel et al., 2024; Lerner et al., 2015; Lopez and Lerner, 2025; Morales and Margolis, 2017; Peña et al., 2025; Poulin et al., 2020), and theoretical models are moving accordingly (Chastain and Redish, 2025; Hayden and Niv, 2021), but see also (Sripada, 2025) for a defense of the common currency hypothesis. Strikingly, a parallel tension has emerged in invertebrate research. In *Drosophila*, dopamine synthesis and receptor signaling are required for olfactory classical conditioning (Kim et al., 2007; Qin et al., 2012; Schwaerzel et al., 2003; Tempel et al., 1984), and activation of specific DAN subsets can substitute for appetitive or aversive unconditioned stimuli (USs) to generate associative memory (Aso et al., 2012, 2010; Burke et al., 2012; Claridge-Chang et al., 2009; Huetteroth et al., 2015; Liu et al., 2012; Yamagata et al., 2015). Many publications describe DAN subpopulations as mediating “reward”, “punishment”, or “reinforcement” (Claridge-Chang et al., 2009; Jovanoski et al., 2023; McCurdy et al., 2021; Musso et al., 2019; Stern et al., 2019; Thoener et al., 2021; Weiglein et al., 2021, 2019), terminology that implicitly suggests a generalized value signal analogous to the “common currency” models proposed for mammals. Comparable roles have been described in larval *Drosophila* (Lesar et al., 2021; Rohwedder et al., 2016; Schleyer et al., 2020, 2013), and dopaminergic activity has been implicated in value-based decision making more broadly (Gorostiza et al., 2016). The anatomical, signaling, and functional parallels between dopaminergic systems in flies and mammals are now sufficiently well-documented to treat them as implementing related organizational principles (Jovanoski and Waddell, 2025). These findings place DANs at the center of valuation circuits in flies as in vertebrates — and they have generated a literature that, implicitly or explicitly, treats dopamine as encoding generalized reward or punishment signals in both.

Yet the evidence from flies, like that from vertebrates, resists such clean interpretation. Protocerebral anterior medial (PAM) dopamine neurons have been described as encoding reward for sucrose, water, courtship and alcohol (Burke et al., 2012; Huetteroth et al., 2015; Lin et al., 2014; Liu et al., 2012; Scaplen et al., 2020; Shen et al., 2023; Yamagata et al., 2015), while protocerebral posterior lateral 1 (PPL1) DANs appear activated by aversive stimuli across modalities, such as bitter taste, electric shock, heat or cold (Aso et al., 2012; Claridge-Chang et al., 2009; Das et al., 2014; Galili et al., 2014; Kirkhart and Scott, 2015; Mao and Davis, 2009; Masek et al., 2015; Riemensperger et al., 2005; Tomchik, 2013; Vogt et al., 2014) — a clean division that mirrors the vertebrate framework. But different dopaminergic populations can substitute for distinct USs depending on motivational state: one subset substitutes for sugar in hungry animals, another for water in thirsty animals (Lin et al., 2014), and another for courtship behavior in males (Shen et al., 2023). Some DAN subsets appear specialized for short- versus long-term memory formation (Huetteroth et al., 2015; Jelen et al., 2023; Yamagata et al., 2015). DANs described as “reward-mediating” in classical conditioning failed to produce equivalent effects in an operant matching task (Rajagopalan et al., 2023), suggesting they may signal expected outcomes rather than reward per se. Recent work shows that the behavioral effects of some DANs are only partially mediated by dopamine release (Mohammad et al., 2024), while neurons previously classified as reward-encoding can drive aversion under some conditions (Lozada-Perdomo et al., 2025). Likewise, while artificial PPL1-γ1pedc neuron activation provides the strongest aversive US substitute (Aso et al., 2012), its activity is also required for artificial octopamine-induced appetitive memory (Burke et al., 2012). Compounding this complexity, broad optogenetic activation of about 40% of the neurons in the PAM cluster — simultaneously recruiting many heterogeneous DAN subtypes — produces a supranormal compound reward signal that drives punishment-resistant, need-indifferent reward-seeking qualitatively unlike the effect of activating any individual DAN subset (Jovanoski et al., 2023), suggesting that the apparent generality of dopaminergic reward signals may depend critically on how many and which neurons are simultaneously activated. The question of whether individual DANs encode generalized value signals or serve more specialized functions is therefore open — in flies as in mammals.

Resolving this contradictory situation requires a direct behavioral test: If specific DANs encoded generalized reward or punishment, animals should consistently seek or avoid their activation regardless of the behavioral context used to reveal that preference. Optogenetic self-stimulation experiments are particularly well-suited for such an experiment. We systematically tested available *Drosophila* DAN subpopulations for their ability to drive consistent appetitive or aversive self-stimulation behavior across a broad range of experimental paradigms: freely moving groups expressing preferences in T-mazes, tethered individuals controlling a platform (“Joystick”) with their legs, and single flies navigating Y-mazes, using optogenetic self-stimulation across multiple light wavelengths. Consistent approach or avoidance across these diverse conditions would support a generalized dopaminergic value signal. Context-dependent effects would argue against it. The breadth of this study — spanning paradigms, preparations, and a decade of systematic experimentation — positions it to provide an unusually direct test of whether dopamine neurons in *Drosophila* implement anything resembling a common currency of value.

## Methods

The project proceeded in three phases. First, we screened 52 dopaminergic driver lines, each expressing the optogenetic channel in a different subset of DANs, in four different behavioral experiments (data collection: 2016-2018). Then we rescreened candidate driver lines in a subset of experiments to test the reproducibility of our screen results (data collection: 2019-2021). Finally, as the last remaining driver line contained DANs in several DAN clusters, we tested lines driving in each of these identified clusters individually (data collection: 2022-2026).

### Screens

The four screens constitute the first phase of this project and data collection was performed in the years 2016-2018. Several key aspects had to be considered when designing a suite of experiments testing the “common currency” hypothesis.

*Operant self-stimulation.* Because most existing work on DAN valence has used classical (Pavlovian) conditioning paradigms — where artificial stimulation is delivered *non-contingently*, meaning that the animal’s behavior during stimulation is irrelevant (e.g. Aso et al., 2012, 2010; Huetteroth et al., 2015; Lin et al., 2014; Liu et al., 2012; Scaplen et al., 2020; Yamagata et al., 2015)— we designed all screen experiments as operant self-stimulation paradigms, in which optogenetic stimulation was contingent on the animal’s own behavior. Of the few existing operant studies, most have included predictive sensory stimuli that the animal learns to associate with optogenetic activation (e.g., Mohammad et al., 2024; Rajagopalan et al., 2023; Stern et al., 2019); such designs introduce classical conditioning processes into what is nominally an operant paradigm (i.e., operant world-learning; Brembs, 2009; Brembs and Plendl, 2008; Colomb and Brembs, 2010). To avoid this confound, we excluded all explicit predictive sensory stimuli, making the animal’s own behavior the sole predictor of stimulation. Only phases in which the fly had active operant control of the light were used for screening; phases in which the light was permanently switched off were excluded from analysis. In other words, we distinguished between operant *activity* and operant *memory/conditioning* which may occur after extended periods of operant activity (Wolf and Heisenberg, 1991).

*Behavioral contexts*. We selected four paradigms that differ systematically in their procedural features, such that they serve as mutual controls. The two T-maze tests groups of freely moving flies; the Y-maze and joystick test individuals. The T-mazes and Y-maze both require coordinated, directed locomotion; the joystick only requires reduced leg flexion/stretch. The joystick immobilizes flies via tethering; all other paradigms allow free movement. Each fly was naive at the start of its experiment and was used in only one paradigm, ensuring that any preference expressed reflects operant exploration rather than prior experience. Because of these built-in procedural contrasts, any consistent behavior across all four paradigms cannot be explained by a single procedural feature — it must reflect something intrinsic to the neurons being activated.

*Optogenetic stimulation parameters.* Stimulation protocols vary widely across the literature, from undisclosed parameters (König et al., 2019) to 10 ms pulses at 50 Hz (McCurdy et al., 2021) to 250 ms pulse duration (Stern et al., 2019) to single pulses exceeding 1.2 seconds at 0.25 Hz (König et al., 2018; Riemensperger et al., 2016). A systematic study found that dopamine release scales with pulse width, frequency, and duration across these ranges (Shin et al., 2020). We used red and yellow light at parameters falling within the effective range established in the literature (see individual paradigm descriptions below for specific values), choosing two wavelengths to allow cross-validation of results across stimulation conditions.

*Controlling for light-evoked behaviors.* CsChrimson is a red-light-sensitive cation channel that depolarizes neurons upon exposure to light peaking at approximately 590 nm (Klapoetke et al., 2014). It was specifically developed to reduce light-induced behavioral artifacts in *Drosophila*, offering better tissue penetration than earlier channelrhodopsins (Klapoetke et al., 2014). Nevertheless, in pilot experiments we found that both red and yellow light at the intensities required for robust CsChrimson activation elicited robust photopreference in wild-type flies in the T-maze (but was without effect in the Joystick). We therefore used blind flies carrying the *NorpA* mutation throughout all experiments to eliminate visual responses to the stimulating light as a confound (see below).

#### T-maze

The T-maze tests the valence of optogenetic DAN activation in groups of freely moving flies making a binary spatial choice. Flies that find optogenetic neuronal activation aversive should accumulate in the dark arm; flies that find it appetitive should accumulate in the illuminated arm. The resulting preference index provides a simple, high-throughput measure of valenced artificial DAN activity across large numbers of animals.

Each T-maze consists of a core unit and three removable tubes, as previously described (Gorostiza et al., 2016): one entrance tube and two choice tubes, one dark and one illuminated (Fig. 1A, B). Flies were loaded into the entrance tube and transferred via an elevator to the choice point, where they could move freely between the two arms for 60 seconds. The illuminated arm delivered optogenetic stimulation contingent on the fly’s position — flies in the lit arm received stimulation, flies in the dark arm did not — giving animals continuous operant control over their level of self-stimulation throughout the choice period. We ran two independent T-maze screens, one with red light (660 nm, 1600 lux, 10 ms pulse width, 20 Hz) and one with yellow light (590 nm, 1000 lux, 10 ms pulse width, 20 Hz), using separate cohorts of flies. LEDs were mounted on a cooling plate to prevent thermal artifacts.

Approximately 30 flies were introduced per run. After loading, flies were held in the elevator for 30 seconds to acclimate before being released into the choice arms. After 60 seconds, the elevator was raised and the flies in each arm and the elevator were counted separately under CO_2_ anesthesia. For the red-light screen, the same groups were tested on two consecutive days with a one-day CO_2_ recovery interval between sessions, allowing assessment of consistency across days. As this measure appeared to be unnecessary, the yellow-light screen was conducted as a single session with double the sample size from the red-light screen; to control for handling variability, scoring was performed blind and in parallel by two independent experimenters.

#### Y-mazes

The Y-maze tests optogenetic valence in individual freely moving flies navigating a three-armed maze, providing a continuous measure of preference at single-animal resolution, as previously described (Werkhoven et al., 2019). Unlike the T-maze, which offers a binary choice between a permanently lit and a permanently dark arm, the Y-maze dynamically tracks each fly’s position and illuminates the designated arm with the optogenetic light whenever the fly enters it, such that the fly’s movement through the maze determines its own stimulation in real time.

The setup consisted of a perspex block (30 × 35 cm) containing 120 individual Y-mazes, allowing simultaneous testing of 120 flies (Fig. 1C), placed in an environmental room at 23° C and 40% humidity. Flies were anesthetized on ice, manually loaded into each Y-Maze with an aspirator and allowed a minimum post-anesthesia recovery period of 20 minutes before tracking. Mazes were constantly backlit with an infrared (invisible for the flies) LED panel and diffuser for homogeneous illumination for the tracking camera. Optogenetic stimulation was delivered via a projector positioned adjacent to the tracking camera, which projected red light (set at [255 0 0] RGB code; 570–720 nm range, maximum intensity at 595nm) onto the designated arm. Fly positions were tracked simultaneously across all 120 mazes using a digital camera equipped with an 850 nm long-pass filter (Table 2) to prevent stimulation light from interfering with tracking (Buchanan et al., 2015; Werkhoven et al., 2019). Projector and camera were calibrated to arm-level spatial precision prior to each experiment (R² ≥ 0.9998 in projector-camera pixel correlation). Stimulation frequency was set to 75 Hz (matching the projector refresh rate) with a 50% duty cycle; tracking was sampled at 37.5 Hz. Fly behavior was recorded and further processed with background subtraction in Matlab (Mathworks). All the Matlab scripts were run under Matlab version 2015a under Windows 7. The projector stimulation patterns were designed with Psychtoolbox-3 toolbox (http://psychtoolbox.org/) with its third-party dependencies and an Nvidia graphic card.

Each fly was tested for 60 minutes. To control for any innate arm preference, each of the three arms served as the optogenetically stimulated arm for 20 consecutive minutes in sequence, such that stimulation was delivered in whichever arm was designated at that time point and the fly happened to occupy. Preference was assessed as the proportion of time spent in the reinforced arm relative to chance.

For example videos of Y-Mazes in operation see (Werkhoven et al., 2019).

#### Joystick

The joystick paradigm tests valence of optogenetic activation in individually tethered flies making a non-locomotor motor decision, providing a measure of operant valence that is entirely independent of spatial navigation or locomotion. Because the fly is tethered and controls a movable platform with its legs rather than walking through a maze, any preference expressed reflects a motor decision unconditioned by locomotor or spatial factors present in the T-maze and Y-maze paradigms.

Flies were tethered by a 0.7 mm monofilament line glued to the thoracic tergite and placed on a laterally movable platform (width: 5mm; Fig. 1D). Leg movements shifted the platform left or right relative to the fly’s longitudinal body axis, and platform position was continuously monitored by a photoelectric detector (analog output −5 to +5 V, sampling rate 33.3 Hz), as described previously (Mariath, 1985; Wolf et al., 1992). Optogenetic stimulation (yellow, 590 nm, 400 lux, 20 Hz, 25 ms pulse width, i.e., 50% duty cycle) was delivered via a light guide directed at the fly’s head and made contingent on platform position: displacement to one designated side activated the light; return past the neutral midpoint extinguished it. The stimulated side was held constant for each individual fly but counterbalanced across flies.

Each fly was tested for a total of 20 minutes, structured as five alternating 4-minute segments: segments 1, 3, and 5 had the light permanently switched off (spontaneous baseline; “test”), while segments 2 and 4 gave the fly full operant control of the stimulation (Fig. 1D, “training”). This alternating design allowed within-animal comparison of spontaneous platform position preference against light-induced preference, with the baseline segments serving as an internal control for any innate side bias. Only the operant control segments were used for screening; the light-off segments were used solely to establish each fly’s spontaneous preference and subtract it from the ‘training’ preference (see “Data analysis” below). Fly line identity was blinded to experimenters throughout, and experiments were conducted by two experimenters in parallel to control for handling variability.

Joystick example video: https://www.youtube.com/watch?v=z2uOIVYrC0o

**Figure 1:**
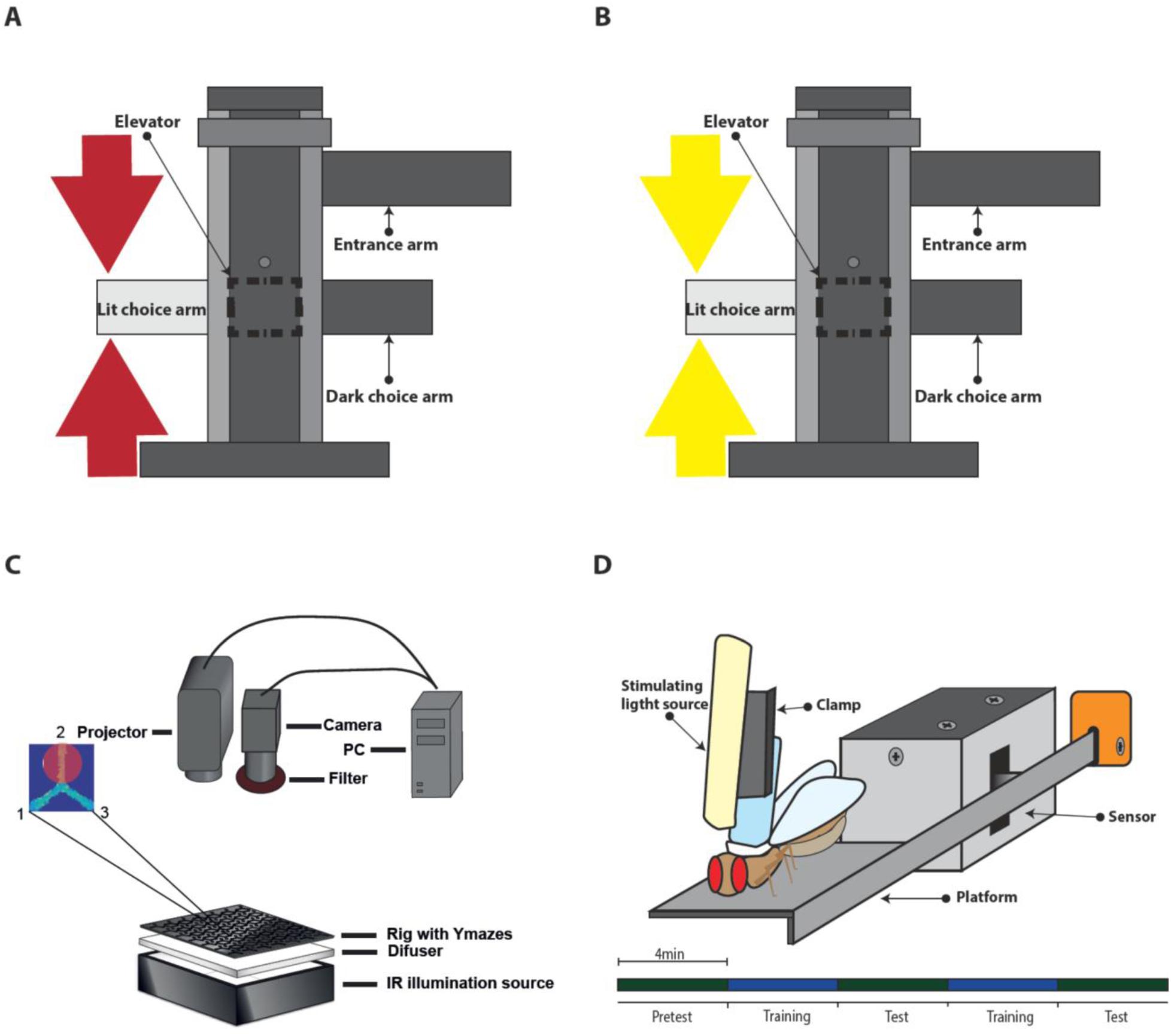
Schematic depiction of the four experimental setups. **A-B**: T-maze schematics. Animals were first loaded into the entrance arm. After acclimatization, they were tapped into the elevator that transported them to the choice point. **A**: Red light was used for optogenetic stimulation (see Materials and Methods for details). **B**: Yellow light was used for optogenetic stimulation (see Material and Methods for details). **C**: Y-mazes schematics. The Y-mazes are IR-illuminated from below and fly behavior is recorded from above. A PC processes the camera image online and switches the projector’s red light for Y-maze arm illumination (enlarged inset) for closed-loop feedback. **D**: Joystick schematics. The tethered flies control the lateral position of the platform with their legs. A sensor records platform position and a PC controls optogenetic yellow light stimulation (via a lightguide) contingent on platform position, which effectively serves as a switch for the light. Bottom: experimental design: ‘training’ denotes periods where the fly controls the light, ‘test’ denotes periods where the light is permanently switched off, see Material and Methods for details.

#### Fly strain selection

In total, we used 52 different fly lines in our four different screens. Of those, 13 were tested in all four screens, another 13 in three screens, 6 lines in two screens and 19 in only one screen (Tab. 1). The selection of driver lines was mostly determined by availability at the time of the experiment. We targeted as many different DAN subpopulations in as many dopamine clusters as was possible at the time. Historically, these were more broadly expressing driver lines, while more specific driver lines were developed later, focussing mostly on DANs with projection in the mushroom bodies (Fig. 2). We added several of these sparse MB-associated DAN lines due to availability and prevalence in the literature. However, as the mushroom bodies themselves are dispensable for spatial preference learning and other forms of learning where olfaction is not required (Mohammad et al., 2024; Putz and Heisenberg, 2002; Wolf et al., 1998), no involvement in operant decision-making was expected or observed. Here, we test the hypothesis that such rewarding or punishing DAN functions generalize to situations beyond a classical US, or whether they merely serve to convey value to an external, likely olfactory stimulus (i.e, in world-learning (Brembs, 2011; Brembs and Plendl, 2008; Colomb and Brembs, 2010)).

**Figure 2:**
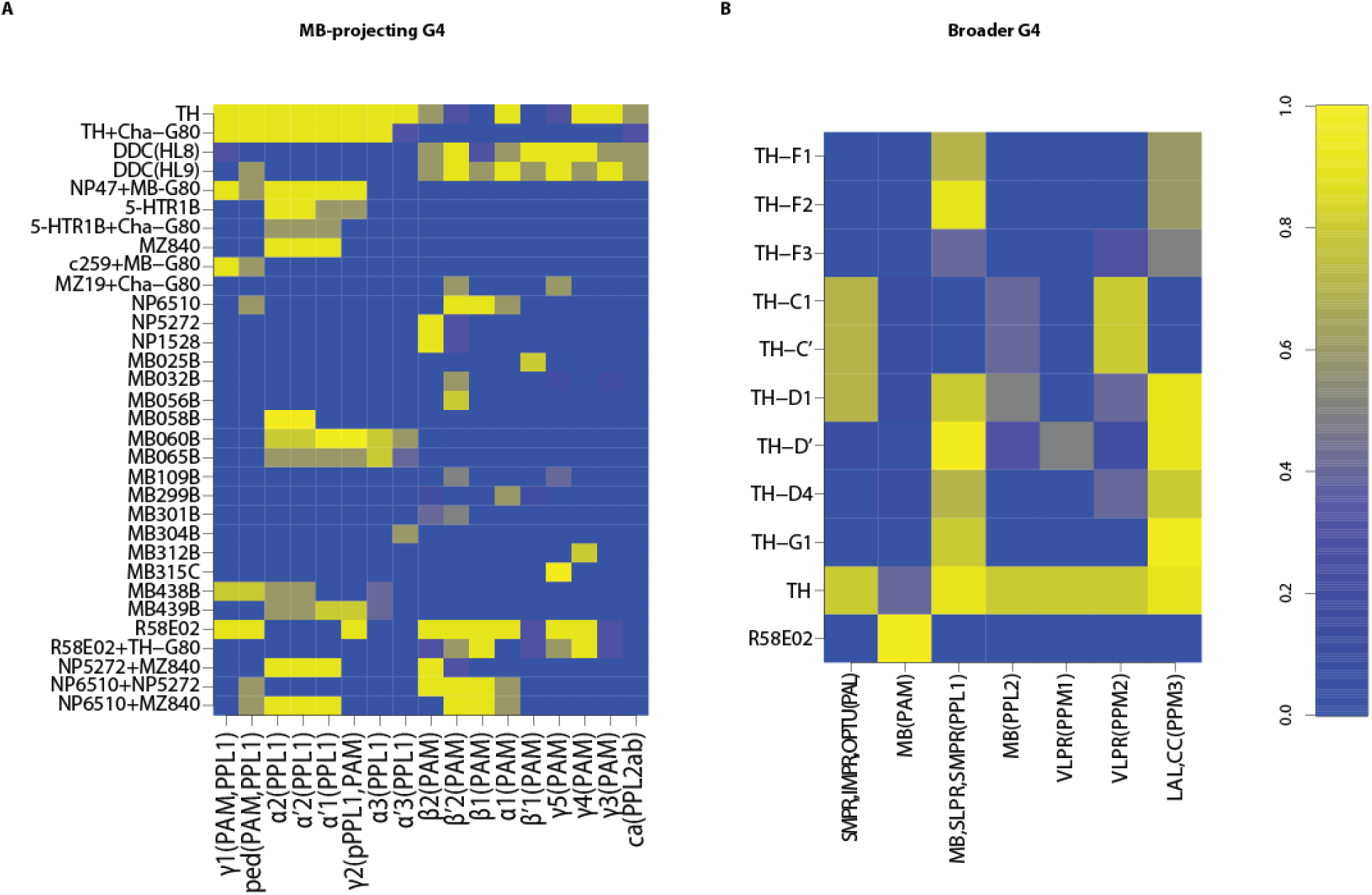
Schematic depiction of DAN driver line expression patterns. Normalized reporter expression intensity from blue (zero) to yellow (one) for each driver line used (vertical axis, see Tab. 1) and their anatomical coverage (horizontal axis; DAN clusters with their projection sites and cell body location, within brackets). **A**: Mushroom-body projecting driver lines obtained from (Aso et al., 2014, 2012, 2010) with their corresponding expression pattern. **B**: Driver lines expressing more broadly than just mushroom-body projecting DANs from (Liu et al., 2012; Pathak et al., 2015). The expression pattern was estimated from (Galili, 2014; Liu et al., 2012; Pathak et al., 2015).

### Rescreening TH-D1 and TH-D’

Rescreens were performed following the four initial screens, testing the two most consistent candidate lines, TH-D1 and TH-D’, with data collection over the years 2019-2021.

#### T-Maze with yellow light

Using identical parameters to those of the original yellow-light T-maze screen, we rescreened the two lines that had performed most consistently across all four paradigms: TH-D1 and TH-D’. We tested a total of seven experimental groups: the positive control Gr28bd;TrpA1 and the two experimental groups TH-D1 and TH-D’ with and without ATR supplementation, and w^1118^;UAS-CsChrimson without ATR to control for potential insertion effects of the effector alone. Data analysis was performed as in the original screen. The values for TH-D1 and TH-D’ were lowest in the T-Maze with yellow light, such that this re-screen provided the most likely route to falsify the hypothesis that these two lines can confer consistent valence over all experiments.

#### Joystick

*Rescreen*. As the T-maze rescreen failed to detect significant effects in the two experimental groups, we not only adjusted our target sample size (see “Statistical Power”, below), but we also tweaked four parameters of the Joystick experiment with the intention to maximize potential effects, while keeping the general self-administration design the same: a) We maximized the light intensity to around twice the levels used in the original screen, i.e., about 900 lux peak intensity, measured at the end of the lightguide. b) To reach the required sample size, we shortened the total experiment duration from 20 to 10 minutes. The periods during which flies controlled the optogenetic light remained identical to the original screen at 8 minutes, but without the intervening test periods. Test periods were now just one minute each, one at the beginning and one at the end of the experiment. c) We increased the required activity levels (“wiggle”, see “Data analysis” below) for inclusion in the analysis from 0.8 to 0.9. d) We performed our statistical analyses on the PIs of the final training period. Due to the large sample size (see “Statistical power” below), we decided against normalizing the training PI to the pretest PI, as in the original screen (see “Data analysis” below). We tested the same seven experimental groups as in the T-maze rescreen (see above).

*Light intensity test*. In order to determine whether the increased light intensity was the cause of the inverted valences of the two experimental lines (see Results), we performed the same experiment as in the rescreen above, but with half the light intensity (i.e., ∼400 lux peak intensity at the end of the light guide) and without the control groups that were not supplemented with ATR. As the valences did not change, we stopped the experiment before the target sample size of 50 was reached (i.e., at N=38).

*Intermittency test*. Original screen and rescreen differed in that the original screen alternated training and test periods, while the rescreen and light intensity test were run with continuous training without interspersed test periods (see above). In order to test whether continuous training was causing the switch in valence of TH-D’, we tested just this line at the light intensity of the initial screen with five alternating periods each of test and training.

### Targeting individual DAN clusters

Following the rescreens, we further dissected the TH-D’ line into different DAN clusters and compared them to the TH-D1 driver line. Data were collected in the years 2022-2026. Fly lines and DAN clusters

Confocal microscopy indicated that TH-D1 and TH-D’ share expression in PPL1, PPM2 and PPM3 DAN clusters. Because we already targeted part of the PPL1 neurons in the original screen, we focused on the PPL1 DANs projecting to the FB body using the driver line SS56699 (Hulse et al., 2021). We addressed PPM2 DANs with TH-C-AD;TH-D-DBD and the PPM3 cluster with TH-FLP-p10;R64H06-Gal4 (Xie et al., 2018). Finally, we investigated the contribution of SIFa neurons using the SIFa-Gal4 driver (Terhzaz et al., 2007), as these neurons were only observed in TH-D1 but not TH-D’ and could potentially explain the different behaviors observed in both driver lines.

#### Joystick with red light

To measure the behavior of flies with CsChrimson expressed using the TH-C-AD;TH-D-DBD line, we equipped the Joystick apparatus with red light. We used the same Philips Lumileds red LEDs as before (see Table 2) and adjusted the light intensity to about 500 lux. Everything else remained as described for the Joystick rescreen above.

#### T-Maze with ten minutes choice time

In addition to narrowing down neuronal populations that drive the behavioral shift in TH-D’ in the Joystick, we investigated whether the shift is consistent across experimental paradigms. For this, we extended the time to choose between the light and dark arm in the T-Maze to ten minutes, but only for strains that showed avoidance in one-minute testing.

#### Dopamine depletion

Dopamine depletion experiments were carried out analogously to previous experiments in our laboratory (Baier et al., 2002). Briefly, laboratory tissue paper (Kimtech Science, Type 7558) was cut to 10 cm x 10 cm and compressed in a glass vial before adding 2 mL of 10 mg/mL 3-iodotyrosine (3IY) dissolved in a 5% sucrose solution.

To prepare the 3IY solution, 2 mL of sucrose solution were added to 20mg aliquots of 3IY powder. For optogenetic experiments, 20 μL of 200 mM ATR were added to the 3IY solution. After vortexing for ∼30 s, undissolved clumps of 3IY were broken down with a micro spatula and vortexed again. The mixture was pipetted onto the tissue paper using a disposable transfer pipette with a large opening, ensuring to also transfer undissolved 3IY residues. Flies were kept for 48 h on aluminum foil-wrapped vials containing both ATR and 3IY before proceeding.

#### Light pre-exposure

We chose five minutes of continuous stimulation to approximate the typical duration of optogenetic light exposure during a Joystick experiment. To expose the flies to the light before a T-Maze experiment, we unwrapped fly vials and placed them between the yellow T-Maze LEDs for the set time. Immediately after pre-exposure, we tested flies in the T-Maze for one-minute, according to the standard protocol, but with only 5 instead of 10 minutes to settle in the loading tube.

### Fly strains, care, genetics and reagents

#### Fly strains used

**Table 1.** Overview of driver lines per paradigm. Also see (Aso et al., 2012) for unlisted stocks. (Co) indicates no all-*trans*-retinal (ATR). * Only one fly was tested.

| Fly line | RRID/FBId | Red T | Yellow T | Y | Joy |
| --- | --- | --- | --- | --- | --- |
| 58E02 | BDSC_41347 | + | + | + | + |
| 5HTR1B | BDSC_27636 | + | + | + | + |
| MZ19 | BDSC_34497 | + | + | + | + |
| NP1528 | DGGR_112679 | + | + | + | + |
| NP0047 | DGGR_112021 | + | + | + | + |
| NP5272 | DGGR_113659 | + | + | + | + |
| NP6510 | DGGR_113956 | + | + | + | + |
| TH-D' | FBtp0083573 | + | + | + | + |
| TH-D1 | FBtp0083568 | + | + | + | + |
| TH-D4 | FBti0151783 | + | + | + | + |
| TH-F2 | FBti0151786 | + | + | + | + |
| TH-F3 | FBti0151787 | + | + | + | + |
| TH-G1 | FBti0151784 | + | + | + | + |
| Gr28bd;TrpA1 |  | - | + | + | + |
| DDC(HL8) |  | - | + | + | + |
| MB025B | BDSC_68299 | - | + | + | + |
| MB056B | BDSC_68276 | - | + | + | + |
| MB065B | BDSC_68281 | - | + | + | + |
| MB109B | BDSC_68261 | - | + | + | + |
| MB304B | BDSC_68367 | - | + | + | +* |
| MB315C | BDSC_68316 | - | + | + | + |
| MB299B | BDSC_68310 | - | + | + | + |
| MZ840 | FBti0004385 | + | + | + | - |
| TH-C1 | FBtp0083567 | + | + | + | - |
| TH-F1 | FBti0151785 | + | + | + | - |
| DDC(HL9) |  | + | + | - | + |
| TH | BDSC_8848 | + | + | - | + |
| MB032B | BDSC_68302 | - | + | + | - |
| MB060B | BDSC_68279 | - | + | + | - |
| MB312B | BDSC_68314 | - | + | - | + |
| TH-C1;TH-F1 |  | - | + | - | + |
| TH-G4;TH-G80 |  | - | + | - | + |
| C259 | FBti0072288 | + | - | - | - |
| GR66A | BDSC_57670 | + | - | - | - |
| GR66A(Co) |  | + | - | - | - |
| TH-C` |  | + | - | - | - |
| GR28bd;TrpA1(Co |  | - | - | + | - |
| 58E02;TH-G80 |  | - | - | + | - |
| 5HTR1B;CHA-G80 |  | - | - | + | - |
| MB058B | BDSC_68278 | - | - | + | - |
| MB301B | BDSC_68311 | - | - | + | - |
| MB438B | BDSC_68326 | - | - | + | - |
| MB439B | BDSC_75961 | - | - | + | - |
| MZ19;CHA-G80 |  | - | - | + | - |
| MZ840N;P6510 |  | - | - | + | - |
| NP0047;CHA-G80 |  | - | - | + | - |
| NP5272;MZ840 |  | - | - | + | - |
| NP6510;NP5272 |  | - | - | + | - |
| TH-G4;CHA-G80 |  | - | - | + | - |
|  |  | T-Maze<br>red | T-Maze<br>yellow | JS<br>red | JS<br>yellow |
| SS56699 | BDSC_87267 | + | + | + | + |
| TH-FLP-p10;R64H06 |  | - | - | - | - |
| SIFa | BDSC_84690 | - | - | + | + |
| TH-C-AD;TH-D-DBD<br>(vs-DAN) |  | + | + | + | + |
|  |  | Dopamine Depletion: T-Maze yellow<br>Pre-Exposure: T-Maze yellow |  |  |  |
| UAS-mCD8:GFP;nsyb-lexA,lexAop-mRFP |  | TH-D1 and TH-D' immunohistochemistry |  |  |  |

#### General fly care

Parental flies were raised on standard cornmeal/molasses medium (Guo et al., 1996) at 25°C and 60% humidity under a 12-hour light/dark cycle. They were flipped daily into fresh vials, to ensure appropriate larval density. Cold anesthesia was used to immobilize offspring flies in the experiments where individual flies were tested.

#### Genetics

To avoid visual cues from the stimulating light that would interfere with the flies’ operant self-stimulation behavior, we genetically blinded flies with a mutated no receptor potential A gene (*norpA^P24^*; RRID:BDSC_9048) (Alawi et al., 1972; Heisenberg, 1971; Ostroy and Pak, 1974; Paj et al., 1976; Pak et al., 1970; Pearn et al., 1996). *norpA* encodes for the phosphatidylinositol-specific phospholipase C that is involved in several sensory pathways (McKay et al., 1995). The mutation abolishes vision completely (Alawi et al., 1972; Heisenberg, 1971; Pak et al., 1970; Pearn et al., 1996). In addition, the *norpA^P24^* allele affects olfactory, gustatory and temperature sensation (Kim et al., 2010; Kwon et al., 2008; Riesgo-Escovar et al., 1995). We crossed the *norpA* mutants with 20xUAS-CsChrimson (RRID:BDSC_82181) flies to create *norpA*;UAS-CsChrimson flies. Because the *norpA* mutation is located on the X-chromosome, virgins from this stock were always crossed with male Gal4 driver lines and the resulting blind male offspring recorded in our experiments.

#### ATR treatment

Male Gal4 drivers and *norpA^P24^;20xUAS-CsChrimson* effector virgin flies were kept overnight on standard cornmeal and molasses food medium (Guo et al., 1996) in darkness at 25°C and 60% humidity for egg laying. One to six days after hatching, groups of approx. 30 male offspring were put in small glass vials with all-*trans*-retinal (ATR) supplementation for two days before testing. For feeding ATR, 15 µl of 200 mM ATR dissolved in ethanol were pipetted onto the surface of 10 ml of food. With these procedures, we ensured that the flies’ first encounter with light was during the experiment. For each setup, pilot experiments determined light intensity and stimulation frequency using a positive control together with a negative control of the same, age-matched genotype, but without ATR in the food. Light parameters were empirically calibrated until the positive control showed a robust effect size relative to the negative control.

#### Positive control lines

In our very first experiments, using the T-maze with red light (Fig. 1A), we used *nor-pA^P24^;Gr66a>CsChrimson* as positive control. This line expresses CsChrimson in bitter taste neurons and their activation mediates avoidance (Marella et al., 2006; Stern et al., 2019). The aversive effects we obtained were of moderate size and so we sought to improve on these controls. In all subsequent experiments, we used *nor-pA^P24^;Gr28bd;TrpA1>CsChrimson* as positive controls. These flies express CsChrimson in heat-sensitive neurons which also mediate aversion. The combination of *Gr28bd* with *TrpA1* drivers showed stronger aversion than *Gr66a* in our experiments and therefore was used for all further experiments. Whereas *Gr28bd* expresses in heat-sensitive sensory neurons in the arista of the antennae (Ni et al., 2013), *TrpA1* expresses in heat-sensitive AC neurons in the central brain (Hamada et al., 2008).

### Optogenetic calibration

Light was measured with a lux meter (Table 2). To confirm that the light spectrum specified in the light source data sheet was correct, the stimulation light was measured with the spectrophotometer at the place where the fly would perceive it (Table 2).

**Table 2.** Equipment used and procurement sources.

| Product | Supplier | URL | article number |
| --- | --- | --- | --- |
| Cosine Corrector for SMA-Connectorized Fiber | Thorlabs | https://www.thorlabs.com/newgrouppage9.cfm?object-group_id=3482&pn=CCS200 | CCSA1 |
| Wide range spectrometer 200-1000 nm | Thorlabs | https://www.thorlabs.com/thorproduct.cfm?partnumber=CCS200#ad-image-0 | CCS200/M |
| Coherent FieldMaxII-TO Laser Power Meter | Edmund Optics | https://www.edmundoptics.de/lasers/laser-measurement/laser-power-meters/coherentreg-fieldmaxii-to-laser-power-meter | #88-411 |
| all trans-Retinal powder, >=98% | Sigma | http://www.sigmaaldrich.com/catalog/product/sigma/r2500?lang=de&region=DE | R2500 |
| Osram Oslon SSL 145 lm yellow 10x10 mm platine LEDs | Osram | https://www.leds.de/osram-oslon-ssl-smd-led-mit-10x10mm-platine-145lm-gelb-69643.html | 69643 |
| Osram Oslon SSL 80 lm blue 10x10 mm platine LEDs | Osram | https://www.leds.de/osram-oslon-ssl-smd-led-mit-10x10mm-platine-80lm-blau-69635.html | 69635 |
| Philips Lumileds red LEDs | Philips | http://de.mouser.com/ProductDetail/Philips-Lumileds/LXM3-PD01/?qs=sGAEpiM-ZZMu4Prknbu83y5qjVwwqxj2nEpXknzdNQZg%3d | LXM3-PD01 |
| Digital Luxmeter 0-400000lx | Voltcraft | https://www.conrad.de/de/luxmeter-voltcraft-lx-1108-bis-400000-lx-kalibriert-nach-werksstandard-ohne-zertifikat-121885.html?WT.mc_id=xplosion&hk=ARX&insert=RA&utm_campaign=xplosiondetailansicht&utm_medium=cpc&utm_source=xplosion&utm_term=121885 | LX-1108 |
| Camera Pointgrey Firefly MV | Pointgrey | https://www.ptgrey.com/firefly-mv-usb2-cameras | FMVU-03MTC-CS |
| Voltcraft DC Power Supply 1,5 to 15 V | Voltcraft | http://www.voltcraft.com/de/labor-netzgerate/ |  |
| Grass Instrument S44G Square Pulse Stimulator | Grass Instruments |  |  |
| Monofilament Line | Cormoran Seacor Shockleader | http://www.cormoran.de/co/de/produkte/angelschn%C3%BCre/seacor_shockleader/5,1,62,63,1,1_products-model.htm?ovs_prdrows2=10&sid=btarcbwsn&stamp=1513450435 |  |
| IR LED panel | Luminous Film | https://www.luminousfilm.com/custom-led-modules.htm |  |

### Immunohistochemistry

#### TH-D1-Gal4 & TH-D’-Gal4

To directly visualize ectopically expressed levels of fluorescent proteins as previously described (Pitman et al., 2011)(Figure 5A-C), we dissected the offspring of UAS-mCD8:GFP;nsyb-lexA,lexAop-mRFP (Weber et al., 2025) virgins and TH-D1-Gal4 or TH-D’-Gal4 males, respectively. In short, brains and ventral nerve cords of 3-5 day old flies were dissected in ice-cold 4% paraformaldehyde solution in phosphate-buffered saline (PBS; 1.86 mM NaH_2_PO_4_, 8.41 mM Na_2_HPO_4_, 175 mM NaCl), and collected in the same fixative on ice for up to 2 hours. Subsequent washing steps (3 x 10 min with PBS containing 0.1% Triton-X (PBST), 2 x 10 min with PBS only) were performed under vacuum and without direct light exposure, followed by mounting in Vectashield Antifade Mounting Medium. For the Multi-Color Flip-Out preparation (MCFO, Figure 5D), we followed the above protocol up to the PBST washing steps, then brains were processed according to the MCFO staining protocol (Nern et al., 2015). In short, brains were preincubated in PBST containing 5% NGS and 5% NDS at 4 °C overnight on a shaker. Under the same conditions, the first primary antiserum was incubated for 2 days, after 5 x 10 min washing in PBST the first secondary antibody was applied for 3 days. Second primary and secondary antisera were incubated similarly for 2 and 3 days, respectively.

Image stacks were taken using a confocal microscope (Zeiss LSM 800) using a 16x objective. See Table 3 for reagents and antibodies.

#### TH-C-AD;TH-D-DBD

Female flies of the driver line were crossed with male UAS-mCD8::GFP flies (RRID: BDSC_5137) and female progeny collected 1–3 days after eclosion. Flies were fixated in 4% paraformaldehyde for 2h at RT before dissecting the brains in PBS. After four washing steps (15 min each) in PBST at RT, nonspecific antibody binding was blocked by incubating brains in 100% normal goat serum (NGS) for 15 min, followed by 10% NGS in PBST for 1h, at RT.

Brains were then incubated overnight at 4°C in the primary antibody solution containing mouse anti-Bruchpilot (1:20), rabbit anti-GFP (1:500), and 3% NGS, in PBST. The following day, brains were washed in PBST at RT four times for 15 minutes before the secondary antibody solution containing goat anti-rabbit AF488 (1:200), goat anti-mouse AF555 (1:200) and 3% NGS in PBST was applied. Secondary AB incubation was overnight at 4°C with samples protected from light exposure.

On the final day, brains were washed in PBST (4x15 minutes, RT) before mounting onto standard microscope slides using Vectashield. Immunohistochemistry samples were imaged using a Leica SP8 confocal laser scanning microscope (RRID: SCR_018169) with a 20x objective and immersion oil. Images were finalized in FiJi (V. 1.54p) and Inkscape (V.1.4). See Table 3 for reagents and antibodies.

**Table 3.** Chemical reagents and antibodies used for immunohistochemistry.

| Chemical | Supplier | Article-/CAS-Number |
| --- | --- | --- |
| Paraformaldehyde | Electron Microscopy Sci-ences | 30525-89-4 |
| Triton-X-100 | Merck | 9036-19-5 |
| Normal goat serum | PAN Biotech | P30-1002 |
| Mouse-Anti-Bruchpilot, nc83 | DSHB |  |
| Rabbit-Anti-GFP | Invitrogen | A-6455 |
| Goat-Anti-Rabbit Alexa Fluor 488 | Mol Probes (MoBiTec) | A-11034 |
| Goat-Anti-Mouse Alexa Fluor 555 | ThermoFisher | A-21127 |
| VECTASHIELD Antifade Mounting Medium | Vector Laboratories | VEC-H-1000 |

### Data analysis and statistics

Since the raw data format was different for each experiment, each data set was analyzed with customized code to obtain a score that allows comparison across setups. Hence, every score ranges from -1 to +1, where negative scores indicate light avoidance, positive scores approach, and close to zero scores indicate no preference for the light.

#### T-mazes

For the T-maze we calculated a Choice Index, CI:

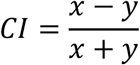

where *x* denotes the number of flies in the bright choice arm and *y* the number of flies in the dark choice arm. The flies caught in the elevator were not considered in the equation as it was unclear if they had not made a decision, were injured and/or impaired in their locomotion, or walking between arms. From the single experiment CIs we calculated the arithmetic mean:

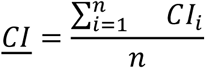

Where *n* denotes the total number of experiments.

#### Y-mazes

For the Y-mazes we calculated the arm occupancy index (OI). Only flies with at least 14 turns/arm changes were used for the analysis and speed was downsampled by window averaging to 3.75 Hz to reduce noise.

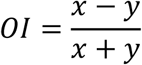

where *x* refers to time spent in the lit arm and *y* refers to the mean occupancy in the dark arms, respectively.

#### Joystick

For the Joystick we calculated the Preference Index, PI, for each of the experimental periods by:

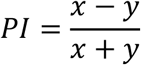

where *x* denotes the number of data points when the light is on and *y* the number of data points in the dark. We normalized the PIs when the flies were controlling the light (‘training’) to the period prior to training (‘pretest’) to only evaluate the change in preference due to the light. In addition, we measured the platform wiggle as a proxy for general activity:

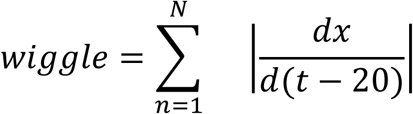

where *x* denotes the time-stamped platform position and *t* the time stamp. The time series was differentiated with a lag of 20 data points to capture the wiggle at a slower time scale that corresponded more closely to the fly behavior. We subtracted the wiggle scores during lights off (test) to when lights were controlled by the animal (training), obtaining a ratio that is positive when flies move more during neuronal activation, and *vice versa*. The wiggle score was used to exclude inactive flies from the data analysis. The wiggle cut-off was set at 0.8 for the screens and 0.9 for the rescreens.

#### Statistical power

Determining appropriate sample sizes for this screen was complicated by the fact that effect sizes for DAN driver lines proved to be substantially smaller than those of our positive controls — something we could not have anticipated before running the first experiment. This meant we could not perform a conventional power analysis before the red-light T-maze screen, and it forced a tradeoff between statistical power, biologically meaningful effect sizes, and practically feasible sample sizes throughout. We therefore approached each paradigm separately.

For the red-light T-maze, no prior power analysis was performed. Based on the effect sizes observed in that screen, we performed a one-tailed t-test comparing positive and negative controls (80% power, α = 0.05) to determine sample sizes for subsequent experiments. This yielded a target of eight runs for the yellow-light T-maze; given our interest in smaller effects suggested by the red-light screen, we set the target at 12. For the T-maze rescreen of candidate lines TH-D1 and TH-D’, we doubled this to 24 to detect the smaller effects observed in those lines with 80% power at α = 0.05. For the joystick, the power analysis yielded a target of 15 flies, which we retained due to practical constraints; for the joystick rescreen, we increased this to 50 to achieve 80% power at α = 0.05 for the approximate preference index of 0.2 observed in the initial screen. For the Y-maze, automated data collection across 120 simultaneous animals generates sufficient observations that even very small effects reach statistical significance; here the relevant question is biological rather than statistical relevance, and no power analysis was performed.

These considerations compelled us to choose an alpha value of 5% (Lakens et al., 2018), despite otherwise routinely aiming for alpha values of 0.5% (Benjamin et al., 2018). Therefore, we supplemented frequentist tests with Bayesian analyses throughout, combining both to arrive at a reliability estimate for each result as indicated in the figure legends.

Code and data availability

#### Code

Most of the analysis and plotting was done in R 2024.04.02 (https://www.R-project.org), except for the data analysis of the Y-mazes, which was done in Matlab 2015a (Mathworks).

Y-Mazes acquisition software: https://github.com/chiser/autoTrackerV2-old-version

Y-Mazes registration scripts: https://github.com/chiser/Registration-Camera-Projector

Joystick instructions sheet with acquisition software (software originally from Reinhard Wolf, Würzburg): https://github.com/chiser/Joystick-acquisition-software

#### Data

All raw data and data evaluation sheets are provided in a single repository. Data and evaluation sheets corresponding to individual figures are organized into separate folders. Data not included in any figure but referenced in the manuscript is available in the ‘Additional data’ folder. https://doi.org/10.5281/zenodo.21722326

## Results

### Screen

#### Robust avoidance with optogenetic feedback

We performed four optogenetic screens on the appetitive or aversive functions of dopamine neuron (DAN) sub-populations in three different operant activity setups. Each screen was designed after a series of pilot experiments had been performed. In these pilot experiments, the experimental details were empirically tuned by testing an aversive driver line (at the time of these experiments, there was no suitable appetitive driver line available) as positive control with and without ATR supplementation. These experiments set the expected effect sizes according to which we estimated our target sample sizes. In each setup, the flies (blinded by the NorpA*^P24^* mutation) were free to choose between a situation in which the neurons selected by the driver line would be stimulated with light and one in which there was no such stimulating light. All four experiments correspond to optogenetic self-stimulation. In the T-maze with red light (Fig. 1A), we used *Gr66a-Gal4* (bitter taste neurons) and in all other setups we used *Gr28bd-Gal4;TrpA1-Gal4* (heat sensing neurons) as positive controls. The positive controls in all four setups show the expected avoidance (negative scores), demonstrating that the basic optogenetic method was working robustly with our chosen driver lines (Fig. 3). Positive control flies tested without ATR supplementation (Fig. 3A,C) showed no light preference or avoidance, confirming that the observed behavioral effects were specifically dependent on CsChrimson activation.

**Figure 3:**
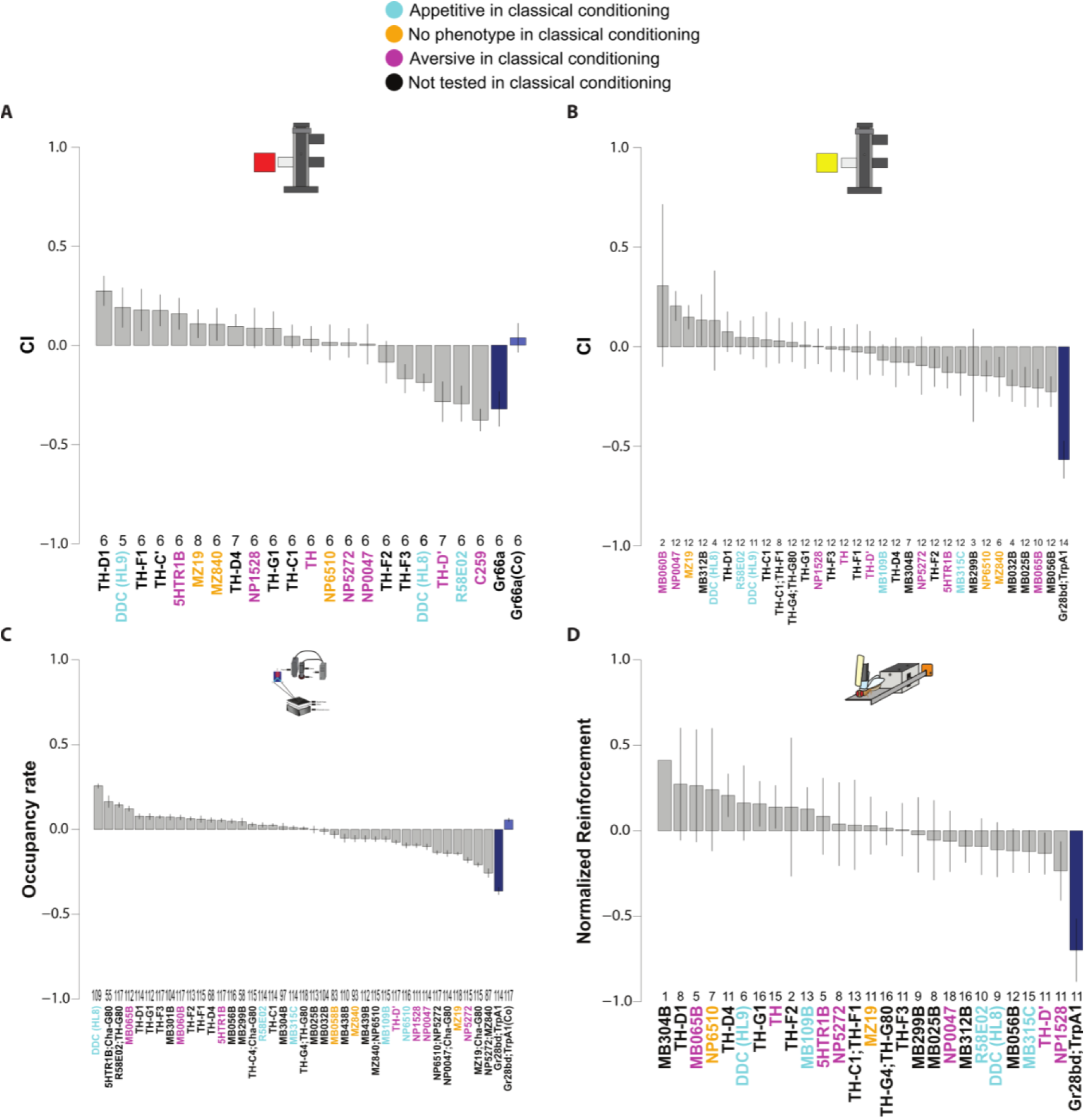
Operant activity screens. **A**: Choice indices from the T-maze screen with red light. **B**: Choice Indices from the T-maze screen with yellow light. **C**: Occupancy Indices from the Y-mazes screen. **D**: Joystick screen Performance Indices. Barplots depict each driver line means for each behavioral score in descending order with error bars depicting the standard error of the mean (SEM). Positive controls fed with and without ATR are colored in dark- and light blue, respectively. The number of experiments for each line is shown above each driver line label on the X-axis. Driver line fonts are color-coded according to classical conditioning phenotypes as shown in the legend above. All lines contained a norpA^P24^ mutation which is omitted for simplicity.

#### DAN activation confers only weak behavioral preferences

Depending on our choice of positive control, the preferences of any of the DAN test lines could have been either larger than the positive control, of the same size, or smaller. None of our positive control lines showed such a strong aversion that any stronger aversion would have been impossible. In fact, when using bitter taste neurons (Gr66a) as positive control, there were two DAN lines with similar avoidance (TH-D’ and R58E02) and one with marginally higher avoidance (C259; Fig. 3A). On the appetitive end, TH-D1-induced light preference was equally strong as Gr66a-induced light avoidance (Fig. 3A). In contrast, in the cases where we used drivers in heat-sensitive neurons as positive controls, the effect sizes were considerably higher than any of the effects observed in the DAN test strains (Fig. 3B-D).

#### Contingent and non-contingent DAN activation confers different valences

Given that their contribution to operant choice is relatively small overall, the question arises if at least the general valence in terms of appetitive, aversive or neutral is conserved, when comparing our results with those reported in the literature for classical conditioning (Aso et al., 2014, 2012, 2010; Liu et al., 2012). Our results suggest that the ability of a DAN subpopulation to replace exafferent appetitive or aversive, respectively, unconditioned stimuli (USs) in classical conditioning experiments does not predict behavior in autogenous, operant experiments: DANs that have been reported to serve as aversive USs can be both aversive or appetitive in operant situations, or provide no discernible preference (Fig. 3, lines marked in red). An illustrative example is line NP0047, which avoids the light in the Y-maze, approaches it in the T-Maze with yellow light and shows no preference in both the joystick and the T-maze with red light. In this case, the same neurons can mediate approach and avoidance depending on the situation. Another notable example is 5-HTR1B which does avoid yellow light in the T-maze but approaches red light in both the T-maze and the Y-maze. Removing cholinergic neurons from the 5-HTR1B patterns using Cha-Gal80 increases the approach of the red light in the Y-maze to the second highest level in this screen. The TH line, which was shown to confer aversive properties to an odor in an operant paradigm (Claridge-Chang et al., 2009), did not show any preference in any of our T-maze screens. It should be noted that this widely used TH line covers only 10% of PAM DANs, excluding many DANs capable of replacing the US in appetitive odor associations (Burke et al., 2012; Huetteroth et al., 2015; Yamagata et al., 2015).

DANs that have been reported to serve as appetitive USs in classical conditioning, can also be appetitive, aversive or neutral (Fig. 3, lines marked in blue). An illustrative example is line DDC(HL8), the line with the most extreme preference for light in the Y-mazes. It avoids the red light in the T-maze and shows weak, inconsistent behavior if the light is yellow in the T-maze or Joystick. Perhaps most notably, lines expressing in the PAM cluster, recently referred to as “traditionally considered to encode reward” (McCurdy et al., 2021), do not seem to mediate much of any valence. For instance, line MB032B promotes avoidance in the T-Maze with yellow light, but is neutral in the Y-Maze. Line MB301B is only weakly appetitive in the Y-Maze, while line MB025B is aversive in the yellow T-Maze, but neutral both in the Joystick and in the Y-Maze. Similarly, MB109B is neutral in the T-Maze with yellow light, very weakly aversive in the Y-Maze, but equally weakly appetitive in the Joystick. Hence, the PAM DANs do not seem to be good candidates for general reward-coding neurons.

Finally, while lines reported to have no effect when used in classical conditioning experiments never showed maximal preference in our screens, some of these lines ended up close to the maximal lines in some of our screens, sometimes aversive, sometimes appetitive (Fig. 3, lines marked orange). For instance, line MZ19 avoided the light in the Y-mazes, but approached it in the T-mazes, while the score was nearly exactly zero in the Joystick screen.

#### Operant DAN function is heterogeneous

Given that the contribution of DAN subpopulations to overall operant performance was small and not consistent with classical conditioning experiments, we tested whether their function is at least consistent between operant experiments. We varied both the type of optogenetic activation and the type of operant paradigm to test how consistently DAN activation would modulate decision-making in the flies. We found that all the parameters we varied influenced DAN function.

##### DAN function varies with optogenetic activation

The T-maze screen was performed with both red and yellow light (Fig. 1A,B). Wavelengths corresponding to yellow light activate CsChrimson more effectively than those corresponding to red light (Klapoetke et al., 2014). However, we did not observe an overall increase in the effect sizes for the DAN lines when tested for their preference in a T-maze with yellow light, compared to one with red light (Fig. 3A,B). If anything, the driver lines mediating aversion of the light decreased somewhat in their effect sizes. While the switch from bitter to heat-sensing neurons for the positive control yielded considerably higher avoidance, absolute effect sizes did not increase for the screened DAN driver lines.

What did change with the switch from red to yellow light was the relative performance of the lines with respect to each other. For instance, the CI of TH-D1 and TH-D’ lines shrunk considerably in yellow light, without changing the sign of their scores. In contrast, line MZ840 changed from appetitive (in red light) to aversive (in yellow light). Overall, the proportion of lines with aversive function increased from 6 out of 21 lines (29%) in the red light T-maze compared to 13 out of 31 lines (42%) in the yellow light T-maze, but this may have been due to a potential bias in the function of the additional lines tested. A direct comparison between the lines tested in both the red light and the yellow light T-Maze reveals no clear relationship between performance in one experiment and performance in the other (Fig. 4B). The lack of correlation between the two experiments suggests that activating CsChrimson more effectively with yellow light did not simply enhance the function the neurons were serving in the T-Maze, rather, it changed their function, even in the same behavioral context of a T-maze.

**Figure 4:**
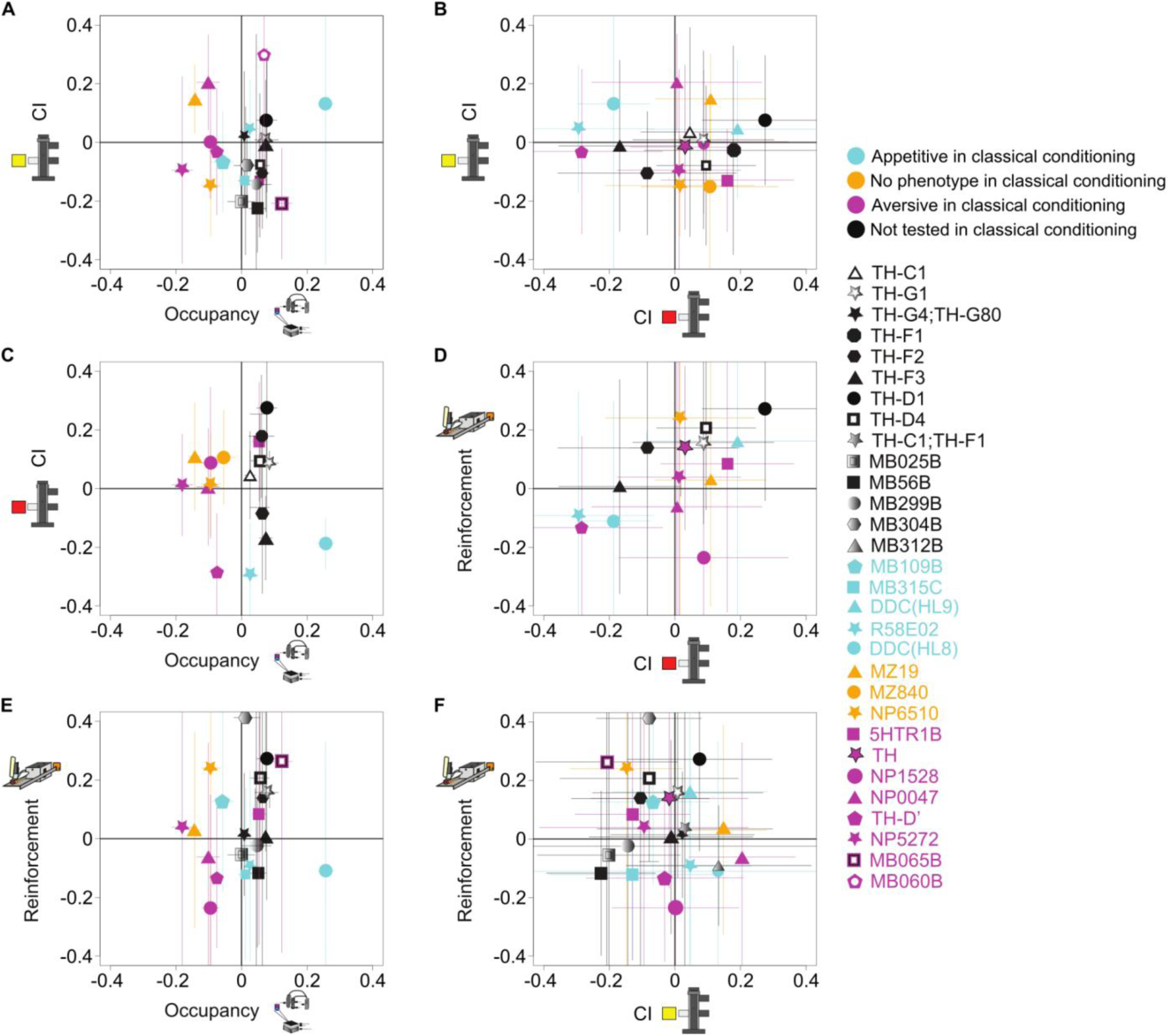
Pairwise comparisons between operant activity scores. Six combinations of biplots projecting two behavioral axes with symbols for their means and bars indicating the SEMs for both axes. Icons depict screen type. Axes ranges were truncated to -0.4 to +0.4 for visualization purposes. Legend on the right, with colors indicating classical learning phenotypes and a corresponding symbol for each driver line.

##### DAN function varies with group vs. individual experiment

The T-maze is an experiment where all the flies in a group are making the same choice at the same time (Fig. 1A,B). The score in these experiments is calculated as the proportion of flies in the illuminated arm, proportional to the total number of flies (Choice Index). To tackle the question of whether an individual will make the same choice when it is alone versus when it is in a group, we performed an analogous forced choice test as in the group-based T-maze in Y-Mazes with individual flies (Fig. 1C). The flies ran continuously in a three-arm Y-maze where one arm was illuminated with red light. We scored the fraction of time spent in the illuminated arm. We chose this measure as it was equivalent to the Choice Index calculated for the T-maze: flies with a longer occupancy of the illuminated arm would have a larger chance of being caught and counted there in the T-maze. This change from group experiment (T-Maze with red light) to single fly experiment (Y-Maze with red light) did not yield any correlation between the scores of the flies in these experiments, not even the most extreme lines were consistent between the two situations (Fig. 4C).

##### DAN function varies with locomotion

In all the experiments described so far, locomotion is a key component of the experiment. In fact, flies that move more slowly in the bright arm of the Y-Maze tend to also have a higher occupancy in this arm (Rohrsen, 2019). As biogenic amines play an outsize role in all kinds of locomotion, we tested whether an operant behavior that is orthogonal to locomotion would yield different functions for our DAN subpopulations. For this test, we placed tethered flies on a manipulandum which they could move perpendicular to their longitudinal body axis using their legs (Joystick, Fig. 1D). Analogous to how a human might move a joystick connected to a computer, the flies could move this platform, on which they were placed, to one side, to stimulate themselves with yellow light, while movement to the other side switched the light off. This transition from locomotion-based to locomotion-independent operant control again yielded a completely new arrangement of lines with regard to their appetitive or aversive function in operant choice (Fig. 4E). In fact, no pairwise comparison between our screens provided a convincing correlation (Fig. 4).

### Three consistent lines

Taking into account the amount of manual labor involved, we had estimated statistical power in two of our setups (T-maze and Joystick) to detect effect sizes at most marginally smaller than those we observed in our pilot experiments with the positive control groups. Only in the Y-Maze screens statistical power was sufficient to detect effect sizes substantially smaller than in our positive controls. The small effects with DAN drivers entail that three of our screens turned out to be considerably underpowered. However, as a whole, all four of our screens might be able to identify the lines that were consistently accompanied by either appetitive or aversive behavior, respectively. Unfortunately, not all lines were tested in all setups due to a variety of technical and historical reasons (see Materials and Methods). Among those 13 that were tested in all four experiments, we searched with the lowest possible criterion of showing the same sign in their scores in all four screens. We found the lines TH-D’, TH-D1 and TH-G1 as the only lines with consistent avoidance/attraction across all four tests. The lines TH-D’ and TH-D1 were the ones with the highest absolute scores; TH-D1 behaved consistently appetitively (i.e., approaching the light) and TH-D’ behaved consistently aversively (i.e., avoiding the light; see Fig. 5 for expression patterns).

**Figure 5:**
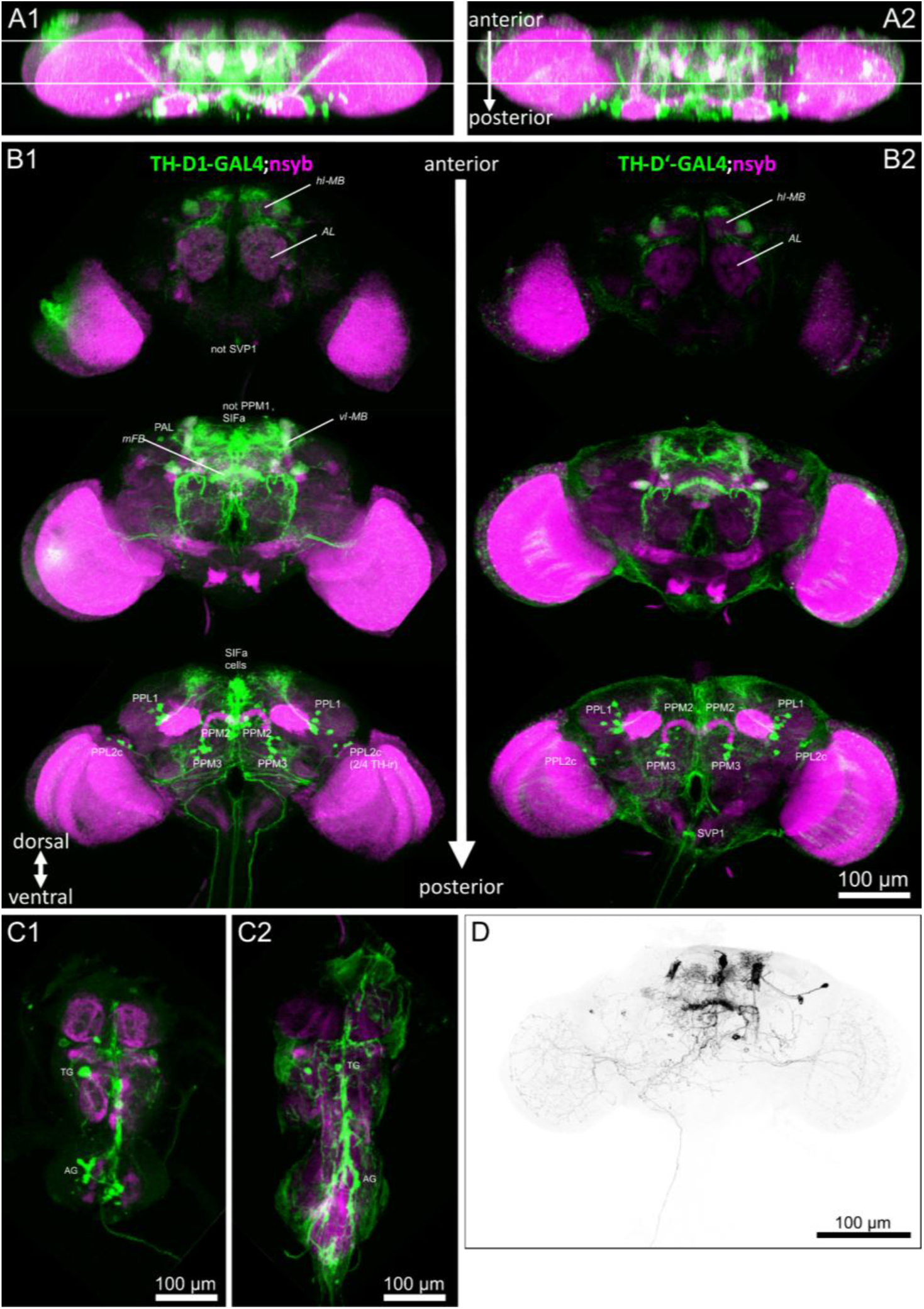
Expression patterns of TH-D1 and TH-D’ driver lines. **A**: Brain, dorsal view. White lines indicate separation of substack projections in B. **B**: Brain, frontal substack projections from anterior to posterior. **C**: Ventral nerve cord. **D**: Stack projection of MCFO-labeled TH-D1-GAL4, revealing the dopaminergic PPL1-α′2α2 and putative PPM3-mFB, and the non-dopaminergic SIFa cell that gives rise to extensive innervation in AL, OL and descending tracts. Green: driver line. Magenta: all neuropil. PPL: paired posterior lateral; PPM: paired posterior medial; mFB: medal fan-shaped body; vFB: ventral fan-shaped body; AL: antennal lobe; vlMB: vertical lobes of the mushroom bodies; hlMB: horizontal lobes of the MB, SIFa: SIFamide neurons.

Interestingly, the two consistent lines express in the same DAN clusters although it was not possible to ascertain whether they labeled exactly the same neurons (Fig. 5). Morphological features at the light microscope-level resolution are often insufficient to distinguish cell types and additional information about synaptic connectivity and genetic expression profiles are necessary to fully define cell identity (Wolff and Rubin, 2018). The cell bodies labeled belonged mostly to DAN clusters PPL1, PPM2 and PPM3 (Landayan et al., 2018; White et al., 2010). At the neuropil level, the two lines labeled specific layers in the fan-shaped body (FB). Staining of the PPM3-vFB (layer 2-4) and the noduli were particularly intense, with TH-D’ FB expression being particularly strong. An additional FB layer from the PPL1-dFB projection was also labeled. The signal in the ellipsoid body (EB) is stronger in TH-D1 whereas the LAL is targeted in both lines (Fig. 5). The mushroom body (MB) vertical lobe and pedunculus projection from the PPL1-y1pedc were clearly visible as well. None of the lines stained the PAM cluster of DANs. The TH-D1 line also labels SIFa neurons in (Fig. 5B1, D), which recently accumulated evidence for co-expressing and co-releasing dopamine (Song et al., 2025).

In order to test the hypothesis that the two most consistent lines in our four screens, TH-D1 and TH-D’, are indeed reliably involved in mediating operant activity, we selected two of our four screens to rescreen these two lines with the appropriate control groups. We estimated our samples to reach 80% statistical power at an alpha level of 0.05. Because of the high alpha value, we supplement these frequentist statistics with a Bayes Factor calculation (see Materials and Methods).

### Rescreen

#### No DAN effects in the T-maze with yellow light

The larger the effects in the first screens, the more likely one would expect their results to reproduce. The two lines show very robust effects in the T-maze with red light and perform at the extreme ends of our distribution of lines (Fig. 3A). While in the Y-mazes, the effect sizes are smaller, the high-powered nature of the screen makes the results there also seem quite reliable (Fig. 3C). Therefore, we sampled from the lowest level of the potential effect size range and began by rescreening TH-D1 and TH-D’ in a T-maze with yellow light. In the original screen, the effects of driving CsChrimson with TH-D1 or TH-D’ were relatively small (Fig. 3B) compared to, e.g., the effects obtained with red light in the same apparatus (Fig. 3A). The sample size in the original screen had been 12 for these two lines and our power analysis indicated that we needed to more than double our target sample size to 25 in order to reliably detect a mean Choice Index of about 0.2. This effect size was chosen for three reasons: a) it was lying just at the edge of the variation in the original screen, b) lower levels of preference would indicate a negligible involvement even if statistically significant and c) practical reasons. At the target sample size of 25 (i.e., about 1,000 flies per group), we could not detect any effects at all in the T-maze with yellow light (Fig. 6A). Compared to their respective control groups, TH-D’ showed a slight tendency to behave aversively and TH-D1 to behave appetitively, as in the original screen, but the effect sizes did not exceed control levels, let alone reach statistical significance.

**Figure 6:**
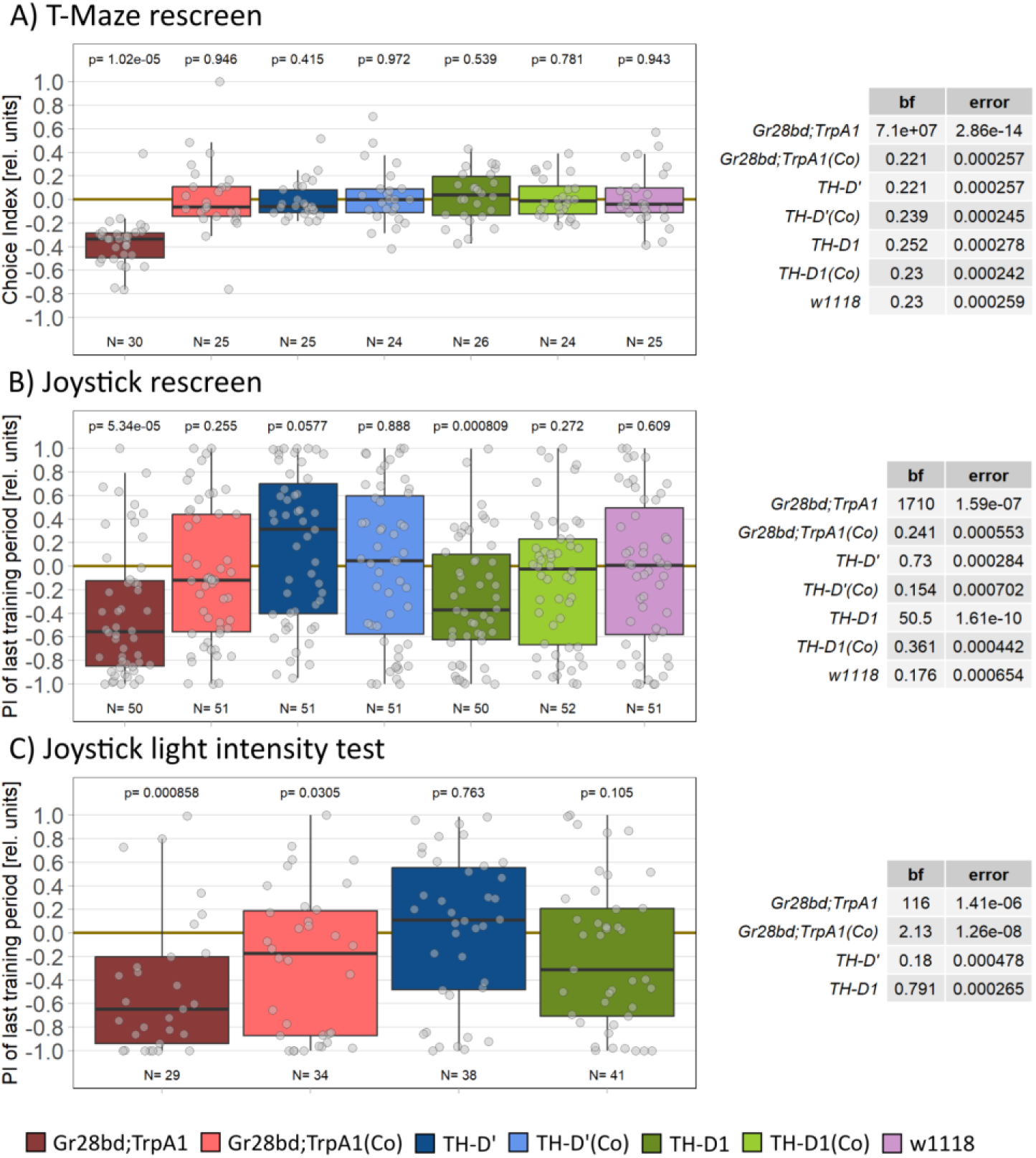
Rescreening TH-D1 and TH-D’ reveals aversive function for TH-D1 in Joystick paradigm. **A**: T-maze rescreen with yellow light. Left: Choice Indices for each line. The control line Gr28bd;TrpA1 shows a statistically significant avoidance of the arm with the stimulating light, whereas all other lines do not show a defined preference. Right: The Bayes Factors of all lines except the control line are smaller than one, providing evidence in support of the null hypothesis of no preference. Together, these data strongly suggest that neither DANs in TH-D1 nor those in TH-D’ support the kind of avoidance behavior that the control flies show. **B**: Joystick rescreen. Left: Performance Indices for each line. Both the control line Gr28bd;TrpA1 and TH-D1 show a statistically significant avoidance of light, whereas line TH-D’ narrowly misses statistical significant preference of the light. All negative control lines do not show any preference, as expected. Right: The Bayes Factors corroborate the frequentist analysis: Gr28bd;TrpA1 and TH-D1 show Bayes Factors larger than one indicating rejection of the null hypothesis of no preference. Together, these data suggest that DANs in TH-D1 can support avoidance in the Joystick experiment. **C**: joystick light intensity test. Left: Performance Indices for each line. Stimulation light intensity was at about 50% of the Joystick rescreen (B) and matched the intensity of the original screen. Main results qualitatively reproduce those in the Joystick rescreen in terms of valence (B). Right: Bayesian analysis as above. Boxplots depict medians (solid lines), quartiles (boxes) and non-outlier ranges (whiskers), with individual data points in gray. Sample sizes below boxplots, p-values for Wilcoxon tests against zero above. Legend below plots indicates color code for groups. Control lines (‘Co’) were not fed ATR, but only solvent.

#### High sample sizes are required for reliable DAN effects

The two most consistent lines ended up towards the extreme ends in our Joystick screen, so we decided to continue with a rescreen in this setup. For the same reasons as above, we also aimed our power analysis for a mean Performance Index of about 0.2, which required a target sample size of 50 in order to reliably detect this effect size. In an attempt to further maximize our chances of detecting any effects, we changed four parameters compared to the original screen (but left everything else identical, see Materials and Methods): a) We maximized the light intensity to around twice the levels used in the original screen. b) We doubled the number of ‘training’ periods to eight (for a total of eight minutes continuous control of the optogenetic light). c) We increased the required activity levels for inclusion in the analysis. d) We performed our statistical analyses on the un-normalized, raw PIs of the final training period. In this rescreen we did detect an effect in the TH-D1 driver line and the valence of the TH-D’ driver line was opposite to that of the TH-D1 line, but the TH-D’ PI did neither reach statistical significance, nor did the Bayes Factor exceed the value of 1 for this group, suggesting that there was no real effect in the TH-D’ line (Fig. 6B). Interestingly, the valence of both lines was opposite to that observed in the four original screens: In this Joystick rescreen, TH-D1 flies avoided the light and the trend in TH-D’ flies was to switch the light on.

As increased light intensity had been shown previously to be able to change valence (Fig. 3), we performed a second experiment with the original light intensity in the Joystick setup as it was used in the screen. However, the valences were unchanged from the previous rescreen (Fig. 6C). Therefore, the increased light intensity in the rescreen was not responsible for the difference in results between screen and rescreen. We conclude, again, that the screen statistical power was too low to reproducibly detect effect sizes as small as those mediated by TH-D’ or TH-D1. Once appropriate sample sizes are reached, even such small effects become reproducible, as can also be seen in a time-dependent valence switch conferred to by TH-D’, below.

#### Operant DAN activation can confer a time-dependent valence switch

A second difference in addition to light intensity compared to the original screen was the temporal structure of the experiment. While in the original screen training periods (where the flies were allowed to switch the light on and off) were interspersed with test periods (where the light was switched permanently off), the rescreen as well as the light intensity test were performed with eight consecutive training periods without intermittent test periods. Therefore, we examined the behavior of the flies during the experiment more closely. Flies with CsChrimson in heat-sensing neurons avoid the light from the start and then tend to slightly decrease their PIs even further (Fig. 7A). TH-D1 flies showed lower, but consistent avoidance throughout the experiment (Fig. 7B). TH-D’ flies, finally, tended to switch the light off at the start of the experiment, only to increase their PIs gradually until the final training period (Fig. 7C). In the light intensity test (see Materials and Methods), both the generally low, negative PI of TH-D1 (available at DOI: 10.5281/zenodo.21722326) as well as the increasing PIs over time in TH-D’ (Fig. 7D) replicated the results from the rescreen. Thus, the increased light intensity in the rescreens did not alter the behavior of these fly strains.

**Figure 7:**
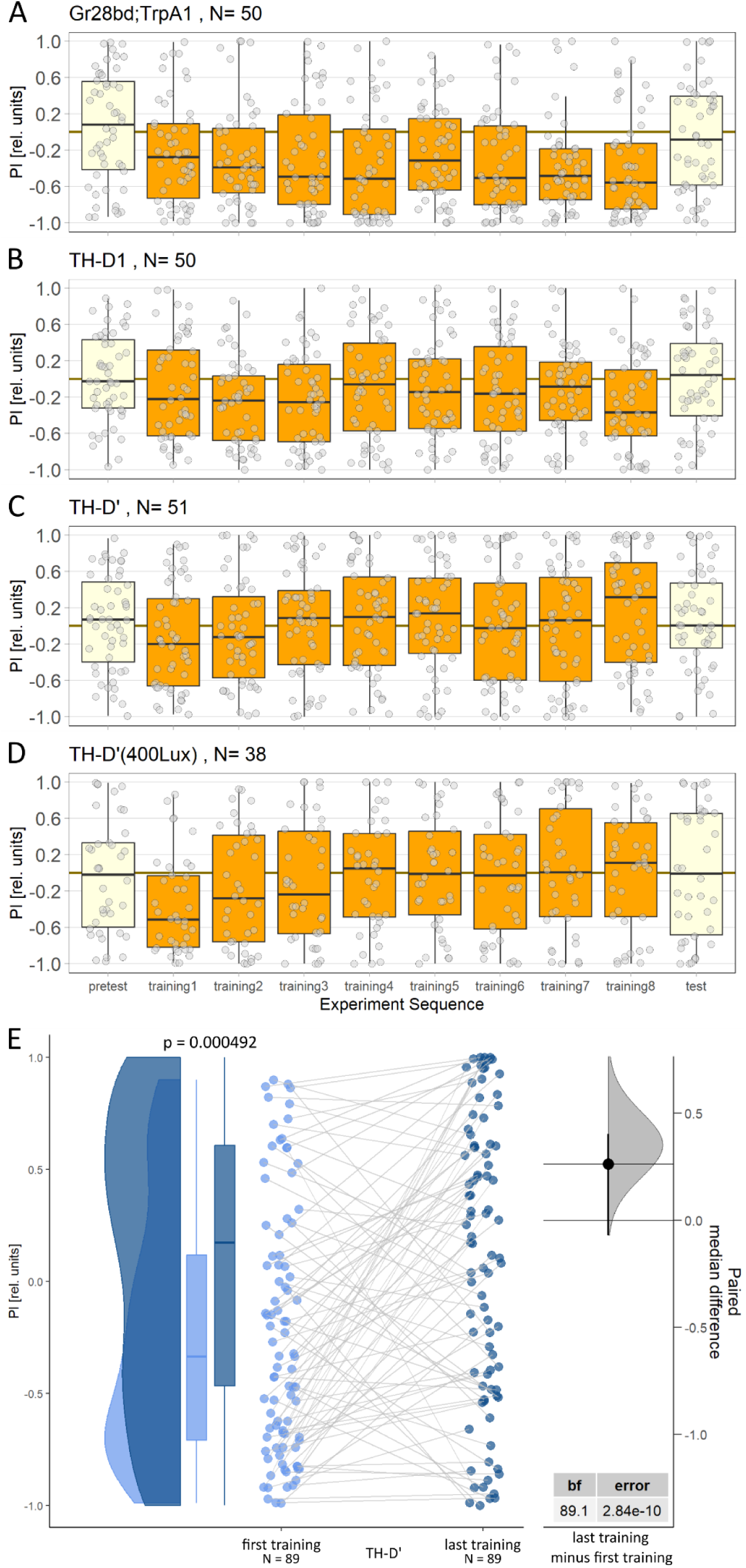
Time course of Joystick experiments reveals a time-dependent valence switch in TH-D’ flies. **A**: Control flies expressing CsChrimson in heat-sensing neurons preferentially keep the optogenetic light switched off throughout the training periods (orange), with little effect on platform position with the light permanently switched off (yellow). **B**: Flies expressing the optogenetic channel in TH-D1 neurons show a markedly weaker but nevertheless consistent preference for light-off during the training periods. **C**: Flies expressing CsChrimson in TH-D’ neurons at 800 lux intensity during the rescreen started their training phase by avoiding the light and finished the last training period with a slight preference for light-on. **D**: The same fly strain as in C tested with 400 lux intensity. The same trend from aversive to indifferent/slight preference can be observed as in C. Time course of box plots depict medians (solid lines), quartiles (boxes) and non-outlier ranges (whiskers), with individual data points in grey. **E**: First and last training differ in TH-D’. Left: Half-violin plot, box/whisker plot and paired individual data plot of the first training period (light blue) and the last one (dark blue). Right: Estimation statistics: Paired median difference of last training (n = 89) minus first training (n = 89) 0.261 [95%CI = -0.0699; 0.401], with Bayes Factor (bf).

Pooling the rescreen (Fig. 7C) with the otherwise identical light intensity test (Fig. 7D) and comparing the first and the last training period reveals the shift in behavior from avoiding the light at the start of the experiment to preferring the light towards the end of training (Fig. 7E). The two training periods are significantly different from each other in a U-test, the BayesFactor suggests a strong effect and the estimation statistics also indicate a shift from aversive to appetitive. In contrast to the last training period, the first training PI is significantly different from zero (Wilcoxon Test, p=0.0003, BayesFactor=99.3). These analyses suggest that for the TH-D’ line, brief encounters with activity in these neurons lead to aversive behaviors, such as those observed in the initial Joystick screen (Fig. 3D). We replicated these findings of the initial screen with a third Joystick experiment where we tested TH-D’ in alternating training and test periods with the lower light intensity from the first screen (see Materials and Methods), where four out of the five training periods indeed showed a negative PI (data available at DOI: 10.5281/zenodo.21722326). Thus, while lines such as NP0047 mediated approach and avoidance depending on the experimental situation, TH-D’ DANs show a time-dependent valence switch.

Taken together, these results show a weakly aversive function for the TH-D1 line in the Joystick experiment (Fig. 6) that does not generalize to other experiments and an initially aversive function for TH-D’ that shifts towards appetitive under prolonged duration of operant control (Fig. 7). The initial aversive function does seem to generalize to other operant experiments (Fig. 2). The switch from aversive to more appetitive function appears to generalize at least across different stimulation intensities (Fig. 7).

### Cluster analysis

Both TH-D1 and TH-D’ are broadly expressing lines, so it is possible that some of their context-specificity arises from the multitude of neurons activated by the optogenetic stimuli and not specifically from a particular group of DANs. With the advent of new, more specific driver lines in the latest phase of this project, it became possible to further dissect the gradual shift from first supporting aversive to later appetitive behaviors in the TH-D’ driver line.

#### Identifying nine bilateral DANs as the source of the valence switch

A prominent anatomical difference between the TH-D’ and TH-D1 lines are the SIFa neurons (Fig. 5). To test if these neurons could account for the differing behavior, we investigated whether isolated activation of these neurons induces any preference in the Joystick. We found no effect (available at DOI: 10.5281/zenodo.21722326), ruling out SIFa neurons as the reason for the difference in TH-D1 and TH-D’ behavior. Next, we tested each of the three DAN clusters that are stained both in TH-D1 and TH-D’ individually. Expressing CsChrimson in PPL1 neurons projecting to the FB did not seem to confer any particular preference for the flies (available at DOI: 10.5281/zenodo.21722326). GFP signal was absent in TH-FLP-p10;R64H06-Gal4>UASmcD8::GFP, therefore we could not test any effect of the PPM3 DANs. The TH-C-AD;TH-D-DBD driver, reported to exclusively target the PPM2 cluster (Xie et al., 2018), revealed additional expression in several PPL1 neurons, as well as optic lobes, KCs, and one suboesophageal zone neuron per hemisphere, putatively SP1 (Fig. 8A). The only overlap between TH-D′ and TH-C-AD;TH-D-DBD, however, are PPL1 and PPM2 neurons. Flies expressing CsChrimson with this driver showed an analogous behavioral shift on the joystick as seen in the TH-D’ driver line, with both red and yellow stimulating light (Fig. 8B). When these flies were tested in the T-Maze for an extended period of time (10 instead of 1 minute choice time) their preference also changed from aversion to attraction with both wavelengths (Fig. 8C). While the effect size varied between the four experiments we conducted, all four of them show the same qualitative behavior that is marked by first avoiding the optogenetic stimulation and then gradually shifting towards preference. This behavior was observed consistently both across experimental setups (Joystick, Fig. 8B; T-Maze, Fig. 8C) and optogenetic wavelengths (red, Fig, 8B1, C1; yellow, Fig. 8B2, C2). Therefore, we abbreviated the name of the TH-C-AD;TH-D-DBD driver line with vs-DAN (valence shifting DANs). The behavior of control flies expressing CsChrimson in heat-sensitive neurons, in contrast, showed no such qualitative variation across experiments or time (Fig. 8C1, C2). The five undepleted vs-DAN 1-minute T-Maze results (Fig. 8C-D) reproduce perfectly the TH-D’ screen results, adding to our confidence that the DANs in the vs-DAN line faithfully represent the DANs responsible for the behavioral effects in TH-D’.

#### The valence switch is dopamine- and contingency-dependent

To test whether simple prolonged exposure to the light — rather than operant control of it — could account for the valence switch, we preexposed flies to approximately five minutes of light, matching the duration typically received by vs-DAN flies during joystick experiments. We selected the T-Maze with yellow light for this test, as it showed the largest effect size. Due to the T-Maze being a group experiment, no yoked control exposure was possible. The pre-exposure reduced T-Maze avoidance in flies expressing CsChrimson in heat-sensing neurons but left behavior unchanged in flies with activated vs-DAN neurons (Fig. 8D1), confirming that the valence switch requires operant contingency rather than mere extended activation. This valence switch is specific to vs-DANs, as, e.g., TH-D1 also started out aversive but did not change valence over the course of the experiment (Fig. 7B). Depleting dopamine with the synthesis inhibitor 3-iodo-tyrosine (3IY,Fig. 8D2) abolished the valence switch in vs-DANflies in the T-Maze entirely, while leaving heat-aversion in Gr28bd;TrpA1 flies in the T-Maze unaffected regardless of dopamine levels (Fig. 8D3). The time-dependent valence switch in vs-DAN flies is therefore both contingency- and dopamine-dependent, whereas Gr28bd;TrpA1-mediated heat-aversion is affected by pre-exposure, but is not dopamine dependent.

**Figure 8:**
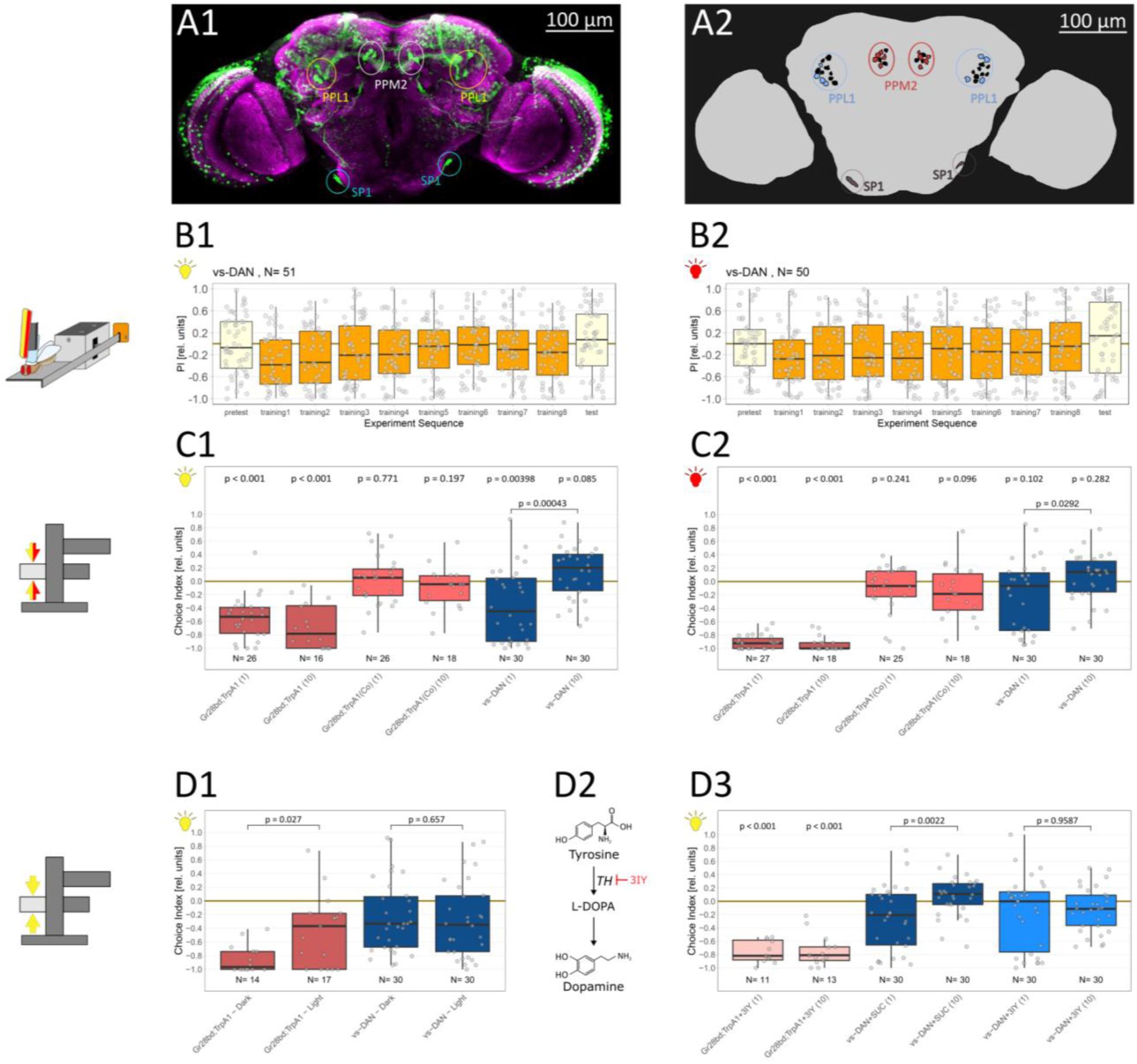
vs-DANs support a valence shift that is contingency- and dopamine-dependent. **A**: Expression pattern of the TH-C-AD;TH-D-DBD (vs-DAN) driver line. Frontal substack projections (A1, green: driver line. magenta: all neuropil) and schematic indicating neurons targeted by the driver (A2, colored: mean number of neurons labeled by the driver, black: total number of cells in the respective DAN cluster); PPL: protocerebral posterior lateral; PPM: protocerebral posterior medial. **B**: TH-C-AD;TH-D-DBD flies show a shift in preference in the Joystick paradigm. In the first minute without optogenetic light stimulation (‘pretest’, light yellow boxes) flies show no preference to move the Joystick platform in a particular position. In the consecutive eight one-minute periods, positioning of the platform to either left or right, respectively, is associated with optogenetic light activation (‘training’, orange boxes). The transgenic flies start out by first avoiding light activation (red light, B1, left; yellow light, B2, right), and transition towards indifference at the end. In both experiments, a slight preference for light activation remains after the light is permanently switched off (‘test’, light yellow boxes). Boxes denote medians and quartiles, whiskers non-outlier range. Grey circles denote individual flies. **C**: TH-C-AD;TH-D-DBD flies show a shift in preference in the T-Maze over time. Different choice times in the T-Maze have little effect on control flies expressing CsChrimson in heat-sensitive neurons, either with (dark red) or without (light red) having been fed ATR (C1 - yellow light, C2 - red light; choice time indicated in parentheses). In contrast, flies expressing CsChrimson in vs-DANs change their interaction with the lit arm from avoidance after one minute to indifference after ten minutes (blue; choice time indicated in parentheses), irrespective of whether the bright arm was illuminated with red (C1) or with yellow light (C2). **D**: The valence shift in TH-C-AD;TH-D-DBD flies is operant and DA-dependent. Pre-exposing flies expressing CsChrimson in heat-sensing neurons to 5 minutes of yellow light slightly reduces their subsequent yellow light-avoidance in a one-minute T-Maze test (D1; red). Flies expressing CsChrimson in vs-PPL1 DANs avoid yellow light irrespective of the pre-exposure (D1; blue). Depleting flies of dopamine by feeding them the DA synthesis inhibitor 3IY (D2) abolishes the vs-DAN-based valence shift in the T-Maze with yellow light (D3; light blue; choice time in parentheses), but leaves heat-sensing-neuron mediated aversion intact (D3; light red, choice time in parentheses). Control vs-DAN-flies show the significant shift from aversive to appetitive after 10 minutes in the T-Maze (D3; dark blue; choice time in parentheses). Boxes denote medians and quartiles, whiskers non-outlier range. Grey circles denote groups of about 30 flies. Numbers above boxes denote p-values for Wilcoxon tests against zero and numbers above brackets T-tests between groups.

## Discussion

Valence perception is the ability to assign subjective value - either positive or negative - to a stimulus or situation (Berridge, 2019; Tye, 2018). We find that both bitter-sensing and heat-sensing neurons can be optogenetically stimulated to replace punishing stimuli and thereby confer negative valence. The explanation for this observation is straightforward: the optogenetic stimulation mimics actual punishing stimuli such as bitter taste and heat. In our operant setups, the behaviors that switch the optogenetic light on lead to positive punishment and the behaviors that switch it off lead to negative reinforcement (escape/relief). The consequence of these two contingencies is the observed robust avoidance behavior in our positive control strains across all behavioral contexts, analogous to classic avoidance behavior in other animals (e.g., Dinsmoor, 2001; Sidman, 1955, 1953). If these sensory neurons were to instruct one set of downstream dopaminergic circuits to represent punishment - for instance via PPL1 neurons previously characterized as mediating aversive stimuli (Aso et al., 2012; Das et al., 2014; Jovanoski and Waddell, 2025; Kirkhart and Scott, 2015; Mao and Davis, 2009; Masek et al., 2015; Riemensperger et al., 2005; Vogt et al., 2014) - and another set for mediating reinforcement - for instance the PAM neurons previously characterized as mediating appetitive stimuli (Burke et al., 2012; Huetteroth et al., 2015; Lin et al., 2014; Liu et al., 2012; Scaplen et al., 2020; Shen et al., 2023; Yamagata et al., 2015) - one aspect of the “common currency hypothesis” would have been corroborated.

### No ‘common currency’ DAN function in flies

If dopamine were universally and critically involved in mediating valence, one would expect dopaminergic neurons (DANs) to be both necessary and sufficient for reward and punishment. To establish necessity, depleting dopamine should severely reduce approach and avoidance behaviors to rewarding or punishing stimuli, respectively. If experimental animals used any behaviors at their disposal to actively seek out activation of DANs mediating appetitive value and avoid activating DANs mediating aversive value, then these DANs would not only be classified as sufficient for computing such value, but the term “common currency” would accurately describe their function.

Our results do not meet any of these expectations. The DAN populations we screened support unexpectedly weak (Fig. 3) and fickle (Fig. 4) approach/avoidance behaviors, compared to our control lines expressing the optogenetic channel in heat sensing neurons. This means that at least the subpopulations we screened do not contribute much to overall approach/avoidance behavior. The results suggest that Dopamine only contributes a minor part to ongoing behavior. Moreover, the negative valence conferred by expressing CsChrimson in these sensory neurons is not dopamine-dependent (Fig. 8), demonstrating that subjective value can be derived without involving dopamine.

### Neuronal substrates of reinforcement

#### Protocerebral anterior medial (PAM) cluster

DANs from the PAM cluster projecting to mushroom body substructures have long been considered to encode reward (Burke et al., 2012; Das et al., 2014; Lewis et al., 2015; Lin et al., 2014; Liu et al., 2012; McCurdy et al., 2021; Yamagata et al., 2015). However, not even the commonly used driver line 58E02 for the PAM cluster showed a positive preference in our test with the highest statistical power, the Y-Mazes (Fig. 3). On the contrary, removing TH-expressing neurons from the 58E02 driver with TH-Gal80 conferred an appetitive valence, suggesting that if there is any valence associated with the neurons targeted by 58E02, they are not part of the PAM cluster. One may ask, why none of the lines expressing in this cluster seem to be relevant for operant activity as tested here. One explanation is that the previous results were gained from classical conditioning experiments with appropriately starved or water-deprived animals, while our flies were satiated. This explanation would be consistent with the activity patterns of PAM DANs during feeding (May et al., 2020). Perhaps PAM DANs are specific to food-related stimuli, rather than mediating general reward? Moreover, operant control has been shown to be independent of mushroom body function across a number of paradigms (Brembs, 2009; Brembs and Wiener, 2006; Putz and Heisenberg, 2002; Siwicki and Ladewski, 2003; Wolf et al., 1998), at least as long as odor processing is not involved. Hence, it may not be surprising that none of the DANs projecting to this neuropil supported consistent preferences in our screens (Fig. 3). As food or water are not the only rewards for flies, none of this should constitute an obstacle for identifying DANs encoding generalized reward. However, while we found DANs that mediated solid preference in each screen, we could not find any DANs that mediated consistent light preference across all screens and held up in our rescreens.

#### Protocerebral posterior lateral (PPL) cluster

Conversely, DANs from the PPL1 cluster, also projecting to the mushroom bodies, have long been considered to contribute to punishment (Aso et al., 2012; Burke et al., 2012; Claridge-Chang et al., 2009; Masek et al., 2015; Riemensperger et al., 2005; Vogt et al., 2014). Several of our lines express in PPL1 neurons. One of them is the line NP0047 with three pairs of PPL1 neurons marked (Plaçais et al., 2012) and it conferred all possible valences in our screens (Fig. 3). Another is line MB438B, only tested in the Y-Mazes showing mild aversion (Fig. 3). Similarly, TH-D’, TH-D1 and vs-DAN all express in some DANs of the PPL1 cluster and at least initially show mild aversion. However, TH-D’ and vs-DAN show a subsequent valence shift (see below). Thus, the evidence about PPL1 DANs does not allow a firm conclusion about this cluster. It is possible that different neurons in this cluster serve different functions.

#### Protocerebral posterior medial (PPM) cluster

Not much is known about the valence conferring properties of DANs in the decisive cluster in the lines TH-D’, TH-D1 and vs-DAN, the PPM2 cluster. Some of the DANs in this cluster have shown a ramping increase in activity during courtship in males (Cazalé-Debat et al., 2024), but these neurons are not necessary for conferring the rewarding properties of courtship (Shen et al., 2023). Whether the time-dependent valence switch reflects an interaction between endogenous ramping activity in PPM2 neurons and exogenous optogenetic activation — or instead arises from plasticity in downstream circuits — to overcome the initial aversion (potentially mediated by PPL1 DAN activity) remains an open question that the connectivity and physiology of this cluster are now well-positioned to address.

In contrast to the dichotomy often thought to exist between separate populations of DANs mediating approach and avoidance, respectively (reviewed in, e.g., Jovanoski and Waddell, 2025), we discovered DANs in the PPM2 and PPL1 clusters (vs-DAN) conferring first aversive and then attractive properties (Figs. 7,8).

#### Pain is so close to pleasure: a dopamine-dependent valence switch

Valence switching is a fundamental adaptive capacity, allowing animals to reassign the positive or negative value of stimuli as conditions change. State-dependent valence switches are well documented: food stimuli become aversive in satiated individuals; in flies, the hungry-to-satiated transition involves opponent neuromodulatory systems (short neuropeptide F and tachykinin) remodeling olfactory processing at the first synapse (Ko et al., 2015); dopamine and octopamine together mediate a photopreference switch when flies lose the ability to fly (Gorostiza et al., 2016); CO2 preference shifts with foraging versus resting state, possibly also via dopamine (Siju et al., 2014; van Breugel et al., 2018; Wasserman et al., 2013); mating state alters odor valence in female *Drosophila* (Boehm et al., 2022; Hussain et al., 2016; Lebreton et al., 2014); and proximity to mating determines whether a looming stimulus triggers escape in males (Cazalé-Debat et al., 2024). Concentration-dependent switches are equally widespread: flies are attracted to low alcohol concentrations but repelled by higher doses (Keesey and Hansson, 2022); salt evokes a concentration-dependent attraction-to-aversion switch in larval and adult *Drosophila* (Dey et al., 2023; McDowell et al., 2022; Russell et al., 2011; Zhang et al., 2013); *C. elegans* approaches several odorants at low doses but avoids them at high doses (Yoshida et al., 2012); and nicotine produces dose-dependent attraction and repulsion in mice (Liu et al., 2022). Similarly, a change in wavelength of the stimulating light, resulting in an altered activation of the optogenetic channel (Klapoetke et al., 2014), could cause a valence switch. Experience-dependent switches include conditioned taste aversion, in which food flavors become aversive after pairing with malaise (e.g., Babin et al., 2014; Nakai et al., 2020; Schafe and Bernstein, 2004; Welzl et al., 2001; Wright et al., 2010; Yoshida et al., 2012; Zhang et al., 2005), or learned phototaxis reversal in several invertebrate species (Crow, 1985a, 1985b; Inoko et al., 1981; Le Bourg and Buecher, 2002; Marchal et al., 2019; Okada et al., 2021; Seugnet et al., 2009). Mere prolonged exposure can also suffice: in mice, extended umami exposure enhances umami preference without affecting bitter avoidance, while extended bitter exposure reduces bitter avoidance without affecting umami preference (Hamada et al., 2023).

The valence switch we observed in TH-D’ and vs-DAN flies is unlike any of the examples above. The animals’ internal state does not change during the ten-minute experiment, ruling out state-dependent switching. The light intensity remains constant throughout, ruling out concentration-dependent effects. The behavior is paired with no other salient stimulus, ruling out experience-dependent conditioning. And critically, mere prolonged pre-exposure to the optogenetic light produces no valence switch (Fig. 8) — demonstrating that operant control of the stimulus, and hence the animal’s own agency, is a necessary condition. The phenomenon itself, however, is not without precedent. Rats and monkeys will repeatedly engage in behaviors that produce nominally aversive stimuli including electric shock and ethanol (Brown and Cunningham, 1981; Cunningham et al., 1993; Cunningham and Niehus, 1997; Kelleher and Morse, 1968; Spealman and Kelleher, 1979), and analogous thrill-seeking behavior has been described in humans (e.g., Gibson et al., 2025; Solomon, 1980; Zuckerman, 2000). The “Opponent Process Theory of Motivation” (Solomon and Corbit, 1974) offers a conceptual framework for understanding such phenomena: stimulus onset and offset are each accompanied by opponent affective processes, and their balance determines overall valence. With repeated exposure, the opponent process strengthens relative to the initial response, gradually shifting the net valence toward the opposite sign. Findings from olfactory conditioning experiments in flies suggest that such opposite valence processes can be mediated by the same DANs releasing different substances (Aso et al., 2019). Applied to our findings, the initial aversive response to vs-DAN activation could reflect the primary process, while repeated operant engagement with the stimulus allows the opponent process to strengthen until approach dominates — a shift that, as we show, requires both dopamine and active behavioral control. Whether this reflects changing properties of the targeted neurons themselves or plasticity in downstream circuits remains an open question, but the finding adds to accumulating evidence that dopaminergic valence signals are more context-sensitive and dynamic than the common currency hypothesis anticipates.

#### Beyond common currency: heterogeneity and context-dependence

Much of the existing fly literature describes DAN subpopulations as mediating ‘reward’ or ‘punishment’ — terminology that implicitly assumes generalization across behavioral contexts. Our results suggest this assumption deserves scrutiny: a neuron that functions as an effective unconditioned stimulus in a classical paradigm, where the animal has no agency, need not function as a reinforcer in situations where the animal does. The weak, fickle, and unstable valence effects we observed across roughly 50 DAN subpopulations converge on a common conclusion: individual DANs do not implement a general-purpose value signal. Instead, our results are more consistent with a view in which DAN function emerges from the combined, heterogeneous activity of many specialized populations — something closer to a combinatorial code than a common currency (Jovanoski and Waddell, 2025). Three features of our data speak to this conclusion, each pointing to a different facet of DAN heterogeneity.

#### Weak DAN effects

While the weakness of optogenetic DAN effects in our operant paradigms violates predictions of the “common currency” hypothesis, it parallels previous findings in both fly and vertebrate research. In operant olfactory and gustatory conditioning, DAN activation produces substantially weaker effects than direct sensory neuron stimulation, a discrepancy attributed to DANs encoding expected rather than actual reward (Jelen et al., 2023; Rajagopalan et al., 2023). This interpretation is consistent with evidence that DAN activity in *Drosophila* tracks stimulus prediction or expectation rather than stimulus delivery (Riemensperger et al., 2005; Vijayan et al., 2022), and with the well-established role of mammalian DANs in computing prediction errors (Gardner et al., 2018; Gershman et al., 2024; Schultz, 2016b; Schultz et al., 1997). A prediction-error account can also explain why DAN activation can produce paradoxically strong effects in classical conditioning: if DAN activity signals that a highly valued event is unexpected, despite this signal being delivered repeatedly with a CS, the postsynaptic circuitry may never fully extinguish its surprise response — potentially explaining the “dysfunctional, unconstrained reward-seeking” observed when large DAN populations are broadly activated as a US (Jovanoski et al., 2023). Apparently, DAN stimulation that is not contingent on the animal’s behavior can recruit processes reminiscent of habit-formation in ways that self-generated DAN stimulation does not.

This discrepancy is reminiscent of other phenomena often observed when studying self-generated stimulation. In general, contingent sensory signals are systematically attenuated relative to non-contingent ones — we cannot tickle ourselves (Bays et al., 2006), we perceive a stable visual world despite repeated saccades (Sommer and Wurtz, 2006), and self-generated and externally imposed stimuli produce distinct brain activation patterns (Matsuzawa et al., 2005). If DAN signals were similarly attenuated when self-generated, their influence on approach and avoidance would be weaker in operant than in classical paradigms. With reafferent sensory feedback, the attenuation may take place via direct inhibition of the sensory pathways (Dallmann et al., 2025; Poulet and Hedwig, 2006), but since DAN activation bypasses sensory afferents altogether, it is not clear how such attenuation would be established. One possible explanation avoiding direct DAN inhibition could be that such operant choices require coordinated activity across multiple neuromodulatory systems rather than a single pathway acting in isolation. Fly photopreference involves opponent contributions from both DANs and octopaminergic neurons, with each system necessary and sufficient for approach toward its respective stimulus (i.e., dopamine for light and octopamine for darkness; (Gorostiza et al., 2016). In mammals, striatal D1/D2 DANs have been proposed to constitute an analogous opponent system (Frank, 2025). If valence signals more generally require such opponent coordination, then activating DANs alone — without simultaneously inhibiting the opposing system — may be inherently insufficient to drive maximal operant effects, regardless of how strongly the DANs themselves are activated. In other words, if a given set of DANs mediated appetitive valence for choosing left, then optogenetic activation would lack the complementary inhibition of the aversive valence associated with choosing right. In contrast, activating heat-sensitive neurons engages both arms of the opponent process downstream, preserving the natural balance of excitation and inhibition. Octopamine lends itself as a candidate for opponent processes in flies as also there, different subsets of neurons can mediate attraction and aversion, respectively, in optogenetic experiments (Claßen and Scholz, 2018) and octopamine is known to be involved in opponent processes with another biogenic amine, its precursor tyramine (Brembs et al., 2007; Roeder, 2005; Roeder et al., 2003; Ryglewski et al., 2017). In mammals, the D1/D2 opponent system is well-described (Frank, 2025), but there are less well-understood interactions between serotonin and dopamine, where both systems need to be involved to accomplish the full behavioral effect (Fischer and Ullsperger, 2017).

Note that the attenuated effect of DAN stimulation on approach/avoidance behavior can also lead to a reduced learned valuation of stimuli associated with reafferent DAN activation, but not necessarily so (Claridge-Chang et al., 2009; Jelen et al., 2023; Mohammad et al., 2024; Rajagopalan et al., 2023). DAN-dependent learning and DAN valence seem to be distinct processes that act through different circuits and signaling systems. Taken together, these results suggest that the distinction between contingent and noncontingent processing may be as important as the identity of the neurons activated — and that generalizing from classical conditioning to operant behavior may be fundamentally inadmissible.

#### Fickle DAN effects

The context-dependence of DAN activation effects in our operant paradigms — where the same subpopulation confers attraction in one paradigm and aversion in another — becomes more intuitive once DAN heterogeneity is fully appreciated. It is already established that different DANs are activated by, and can substitute for, different appetitive stimuli: one subset signals progress toward feeding, another toward drinking, another toward copulation, etc. (Cazalé-Debat et al., 2024; Jovanoski and Waddell, 2025; Lin et al., 2014; Shen et al., 2023), suggesting that the expectation of different future events is mediated by different DANs. From that perspective, each DAN population can be understood as a reporter for a specific goal. In our experiments, the only contingency available to the animal was the optogenetic stimulus itself, and flies are known to behave in ways that reflect the consequences they expect their actions to have (Heisenberg, 2015, 2014). If the same DANs signal different outcomes in different behavioral contexts, this may be because the goals animals pursue differ across contexts — a tethered fly and a freely navigating fly are not trying to accomplish the same thing. For instance, if a tethered fly had managed to free itself from the tether, this event would carry positive valence and activate a set of DANs specific for this goal in this context. Activity in the same set of DANs in a freely navigating fly, on the other hand, may mediate the negative valence of having entered an arm in a maze that does not lead to escape. Activating these same DANs would then carry different values depending on which goal is currently active, just as the same reward circuit encodes different outcomes depending on the animal’s physiological state. We therefore hypothesize that context-dependence of DAN function may reflect goal-dependence rather than ambiguity in the dopaminergic signal itself. There is evidence from birds that this may be a general organizing principle of DAN function (Roeser et al., 2023).

#### Unstable DAN effects

How can the same DANs drive aversion initially and approach minutes later (Figs. 7, 8)? One tempting possibility is that the valence switch reflects learning rather than a property of the neurons themselves. Rats and flies alike prefer situations where they can control or predict (even aversive) stimuli over those where they cannot, even when the stimuli are otherwise identical (Badia et al., 1979; Heisenberg et al., 2001). Relief learning — where actions or stimuli that terminate an aversive stimulus become positively reinforced — provides a potential mechanism (Dinsmoor, 2001; Gerber et al., 2014; Ilango et al., 2012; Mohammadi et al., 2014), and flies are capable of exactly this form of learning (Gerber et al., 2014; McCurdy et al., 2021; Niewalda et al., 2015; Yarali et al., 2009; Yarali and Gerber, 2010). “Opponent Process Theory” offers a complementary account at the systems level (Solomon and Corbit, 1974). Together, these frameworks suggest a unified hypothesis: initial DAN activation, in the absence of learned contingencies, signals lack of control and is experienced as aversive; repeated activation establishes a predictable contingency, and the same signal now indicates that the animal controls its environment, driving approach. Consistent with this, non-contingent vs-DAN stimulation — where no behavioral contingency exists — does not generate any shift in valence (Fig. 8). The observation that TH-D1 neurons drive stable avoidance without a valence switch (Fig. 7B) further suggests that the instability is not a general property of aversive DANs but something specific to vs-DAN, whose detailed circuitry and physiology may therefore be worth investigating as a substrate for controllability-dependent valence encoding. These results reinforce the important distinction between contingent and non-contingent stimulation.

### Limitations

The most significant limitation is coverage. Only 13 of nearly 50 lines were tested across all four paradigms, and while no strong candidates emerged even among the 29 lines tested in two or more screens, we cannot exclude the possibility that an untested combination of DANs would produce consistent, robust operant preference. What we can say is that across the subpopulations we did test, no single line approached the behavioral magnitude of our sensory control strains in any consistent direction. This suggests that the dopaminergic system, at the resolution of individual subpopulations and across the range of operant contexts we examined, does not behave like a system built around a common currency of value. It behaves like a system whose function is distributed, context-sensitive, and inseparable from the goals and expectations of the animal in which it operates. The one exception — the vs-DANs, whose dopamine-dependent valence switch appears tied to the animal’s operant control of its environment — represents precisely the kind of cellular-resolution entry point that connectomic and physiological approaches in *Drosophila* are now positioned to exploit.

There are technical reasons to interpret our results with some caution. Our stimulation parameters — optimized for sensory control neurons — may not be equally effective for non-sensory DANs, whose input-output properties are unknown. Our regimes fall within the effective range found in the literature for DAN activations (see Materials and Methods), but may not be optimal for every subpopulation of DANs. It is impossible to experimentally titrate light stimulation in terms of frequency, duration and intensity, until it is optimal for each chosen population of neurons. That stimulation regimes can in principle affect preference has also been observed in a two-choice optogenetic paradigm over long timescales driving activity in octopaminergic neurons (Claßen and Scholz, 2018). Activating DANs with different intensities and optogenetic effectors in classical conditioning experiments with *Drosophila* larvae can affect the strength of the teaching signal (Toshima et al., 2025). More recently it was observed that modulatory neurons can release different substances at different levels of optogenetic activation, leading to a switch in valence (Savaş et al., 2026). It is also possible that appetitive DAN function in some lines is state-dependent — hunger or thirst gating the signal — though state-independent appetitive DANs exist and should in principle have been detectable in our screens (Lin et al., 2014; Shen et al., 2023).

It is also somewhat unexpected that the avoidance of optogenetic heat is independent of dopamine, as DANs have been associated with temperature preference before (Galili et al., 2014; Tomchik, 2013). At the same time, when heat is used to induce place learning in walking flies, both dopamine and octopamine are dispensable, while serotonin is required (Sitaraman et al., 2010, 2008). The precise function of dopamine in processing temperature sensations in *Drosophila* does seem to require further research.

## Acknowledgments

We are grateful to our dedicated undergraduate students Aida Kumpf, Kader Semiz, Ferruh Aydin, Gaia Bianchini, Avani Koparkar, Amanda Torres, Saurabh Bedi and Naman Agrawal, for their technical assistance in data collection. We are indebted to Hiromu Tanimoto for his generous sharing of fly stocks, as well as to Andreas Thum for sharing the SIFa-Gal4 line. Funding was provided by DFG grant BR 1892/16-1 to BB and HU 2474/1-3 (Project Nr. 371431309) to WH. None of this work would have been possible without the assistance of the electronics and precision mechanics workshops of the University of Regensburg. As non-native speakers, we used different LLMs to improve the language of this manuscript; the authors remain responsible for content and language.

*CRediT*: Conceptualization: CR, BD, BB; Data curation: DD, CR, BB; Formal Analysis: DD, CR, WH, BB; Funding acquisition: CR, BD, WH, BB; Investigation: DD, CR, WH, BB; Methodology: DD, CR, BD, WH, BB; Project administration: BB; Resources: BD, BB; Software: DD, CR, BD, BB; Supervision: BD, BB; Validation: BB; Visualization: DD, CR, WH, BB; Writing – original draft: CR; Writing – review & editing: DD, CR, BD, WH, BB.

## Notes

https://doi.org/10.5281/zenodo.21722327

### Competing Interest Statement

The authors have declared no competing interest.

### Summary of Updates

New experiments addressing reviewer suggestions for more specific driver liens and dopamine involvement. Completely re-written manuscript, Additional conclusions added to old conclusions.

